# Stress-response regulators tune enterobacterial common antigen synthesis to support acid and reactive nitrogen species stress resistance

**DOI:** 10.64898/2026.10.23.761472

**Authors:** Haley C. Hill, Angela M. Mitchell

## Abstract

Enteric bacteria must survive envelope-damaging environmental conditions in both host and environmental niches (e.g., acidic environments, reactive oxygen and nitrogen species). The Gram-negative outer membrane (OM) is central to the cell’s stress resistance, acting as a permeability barrier to antimicrobials and as an adaptable structure remodeled in response to environmental changes. *Enterobacterales* (*e.g., Escherichia coli*, *Salmonella enterica*, *Klebsiella pneumoniae*) produce an invariant glycan, enterobacterial common antigen (ECA), which is localized in the OM and periplasm and contributes to the OM permeability barrier. Yet, how ECA biosynthesis is controlled and the conditions under which ECA is important remain poorly defined. Therefore, we investigated transcriptional regulation of the ECA biosynthesis *wec* operon in *E. coli* K-12. We assayed *wec* operon promoter activity across deletions of 36 bioinformatically identified potential regulator genes and identified four for which deletion significantly altered *wec* promoter activity. Of these, two repressors, GadX, a regulator of the glutamate-dependent acid stress response system, and MetJ, the master regulator of methionine biosynthesis and a reactive nitrogen species (RNS) stress regulator, bound to the *wec* promoter *in vitro*, consistent with direct transcriptional regulation. In accordance with a link between ECA and these regulators, *wec* promoter activity increased under acidic pH and under RNS stress, and loss of ECA impaired growth under both conditions and fitness in acidic pH. Together, these findings reveal that biosynthesis of a conserved OM glycan is embedded within dedicated environmental stress regulons and identify previously unrecognized roles for ECA in bacterial stress adaptation.

**Importance:** Enteric bacteria rely on their cell envelope to survive chemical and physical stresses in changing environments. Enterobacterial common antigen (ECA), an invariant envelope glycan produced by enteric bacteria (*Enterobacterales*), plays a role in the outer membrane (OM) permeability barrier that excludes detergents and antibiotics. Despite its conservation, the cellular function of ECA, and the conditions that require it, remain elusive. Here, we define the transcriptional regulation of ECA biosynthesis and use it to identify environmental conditions in which ECA function is important for cellular growth. We identified regulators of two stress responses (GadX, an acid stress regulator, and MetJ, a reactive nitrogen species stress regulator) as direct repressors of ECA biosynthesis and show ECA promotes growth under both acid stress and reactive nitrogen species stress. These results indicate that a conserved envelope glycan is integrated into stress-response networks and functions broadly in adaptation to environmental stress.

## Introduction

The cell envelope is the frontline of bacterial defense from hostile and changing environmental conditions. Gram-negative bacteria, such as those in the order *Enterobacterales* (e.g., *Escherichia* spp., *Salmonella* spp., *Yersinia* spp., *Shigella* spp.) possess a multilayered cell envelope composed of an inner membrane, a thin peptidoglycan layer located in the periplasm, and an outer membrane (OM), the outermost barrier in the envelope. The OM is crucial to the cell’s resilience to environmental stress, acting as a permeability barrier that excludes antibiotics and detergents (1, 2), a barrier to physical and osmotic stress (2–6), and as an adaptable structure that can be remodeled in response to environmental cues (7–10). The OM is asymmetric: its inner leaflet facing the periplasm consists of phospholipids and its outer leaflet is composed predominantly of lipopolysaccharide (LPS). *Enterobacterales* also produce an invariant surface carbohydrate, enterobacterial common antigen (ECA), which can be present at the OM and in the periplasm (11–13). First described in 1962 by Calvin Kunin and colleagues (14), ECA has been studied for decades; however, its functions have only recently begun to be resolved (15).

ECA has been linked to OM integrity and intrinsic resistance to antibiotics and detergents. Loss of ECA causes increased sensitivity to detergents including bile salts (16). In addition, ECA has been shown to have genetic interactions with YhdP, a protein that contributes to phospholipid transport between the inner membrane and OM (17). In the absence of YhdP, cells become sensitive to vancomycin and the detergent SDS, and this sensitivity is suppressed by loss of ECA (16, 18). Together, these results demonstrated that ECA maintains the OM permeability barrier, and that it can take on aberrant activity that damages the OM permeability barrier in the absence of YhdP.

Beyond envelope permeability, ECA has also been implicated in the pathogenesis of *Enterobacterales*. In *Salmonella enterica* serovar Typhimurium, mutants unable to synthesize ECA show reduced virulence and heightened susceptibility to bile salts (19). High-throughput infection studies further indicate that ECA promotes *Proteus mirabilis* virulence during polymicrobial urinary tract infection (20), enhances recurrent urinary tract infection caused by uropathogenic *Escherichia coli* (21), and supports *Klebsiella pneumoniae* bloodstream infection in mice (22), as well as *Enterobacter cloacae* infection in a wax worm model (23). Together, these findings indicate that ECA supports survival of diverse *Enterobacterales* across host environments. Here, we test the hypothesis that, if ECA contributes to fitness under specific stresses, its biosynthesis might be dynamically controlled by stress-responsive regulatory systems that adjust ECA production to the specific environment.

Untangling further details of ECA function and its interactions with the cell’s environment is made complicated by the sharing of biosynthetic machinery between ECA and other surface glycoconjugates (24, 25). Specifically, biosynthesis of ECA, LPS, peptidoglycan, and O antigen and colanic acid if produced all compete for the same limited pool of undecaprenyl-phosphate (Und-P) the isoprenoid carrier (24) and for some amino sugar precursors (26). Therefore, disrupting ECA biosynthesis can alter flux through other essential Und-P-dependent pathways, complicating interpretation of phenotypes attributed specifically to ECA loss (11, 13, 27). Consequently, different approaches are needed to elucidate cellular ECA functions.

ECA biosynthesis, recently reviewed in detail (15, 28), synthesizes a polymer of five to seven repeating trisaccharide units (16, 29) composed of the amino sugars N-acetylglucosamine (GlcNAc), N-acetyl-D-mannosaminuronic acid (ManNAcA), and 4-acetamido-4,6-dideoxy-D-galactose (Fuc4NAc) (30–32) on Und-P at the inner membrane (30, 33–36). The polymerized chains are transferred to either diacylglycerol phosphate (DAG-P) to form ECA_PG_ or to LPS to form ECA_LPS_ (12, 37, 38) and these molecules are transported across the periplasm to the OM for surface display. A third form of ECA, ECA_CYC_, is cyclized and retained within the periplasm (12, 29, 39, 40). The mechanism of ECA_CYC_ cyclization remains uncharacterized but requires the ECA chain-length regulator WzzE (39). Both ECA_CYC_ and ECA_PG_ play roles in maintaining the OM permeability barrier in *Escherichia coli* K-12 (16).

The genes responsible for synthesis of the ECA amino sugars (27, 40), their loading onto Und-P (41), translocation across the inner membrane (42), polymerization (39), and chain length regulation (43) and cyclization of ECA are encoded in the *wec* operon. Mutations in this operon can inhibit ECA synthesis entirely or affect ECA chain length or abundance (15, 27, 30, 39, 43). Of particular interest, deletion of *wecA* (*rfe*), the first gene in the operon, abolishes production of all three ECA forms (16, 30, 40), while deletion of *wzzE*, prevents formation of ECA_CYC_ and causes linear ECA to be made with unregulated chain length (29, 39, 43). Examining the regulators and environmental conditions that affect *wec* operon expression and ECA abundance provides a possible avenue for identifying conditions in which ECA function is important for cellular homeostasis.

Several regulators of ECA biosynthesis have been reported in recent years (reviewed in (28)). For example, PdeL, a cyclic-di-GMP phosphodiesterase, was identified as the first confirmed transcriptional regulator of the *wec* operon (44), and a post-transcriptional pathway regulating ECA abundance was subsequently discovered (45). To further define the regulatory landscape of ECA biosynthesis, we sought to identify further transcriptional regulators of the *wec* operon in *E. coli* K-12 and to determine whether ECA function is relevant to the environmental conditions in which these regulators are active. After evaluating 36 bioinformatically identified candidate regulators, we identified four genes, *gadX*, *metJ*, *nagC*, and *phoB*, for which deletion altered *wec* promoter (P*_wec_*) activity. Of these, only deletion of *metJ* and *gadX* produced corresponding significant changes in ECA abundance, while Δ*nagC* and Δ*phoB* caused changes to ECA levels contrary to their transcriptional effect. We therefore focused subsequent mechanistic characterization on MetJ and GadX. Consistent with GadX’s established function in acid resistance (46, 47), decreased pH increased P*_wec_* activity and cells lacking ECA had a growth defect in acidic media. MetJ plays a role in resistance to reactive nitrogen species (RNS) stress. We found that S-nitrosoglutathione (GSNO), a bioavailable nitric oxide donor, induced P*_wec_* activity and that mutants lacking all ECA (Δ*wecA*) or ECA_CYC_ (Δ*wzzE*) had growth defect under GSNO treatment. Together, these findings establish MetJ and GadX as regulators of ECA biosynthesis and link ECA production to two distinct cellular stress response pathways — acid resistance and RNS stress.

## Results

### Several potential regulators affect P*_wec_* activity

To determine whether our previously constructed P*_wec_* transcriptional reporter (45) could detect changes in P*_wec_* activity due to altered transcriptional regulation, we measured P*_wec_* activity in a Δ*pdeL* strain using a reporter plasmid (**Fig. S1**) containing the *wec* promoter region (P*_wec_*) driving the luciferase operon, *luxABCDE* (pP*_wec_*::*lux*). PdeL, a cyclic di-GMP-phosphodiesterase and transcriptional regulator, was recently demonstrated to directly repress expression of the *wec* operon (48). The effect of PdeL on P*_wec_* activity was measured in cells grown in minimal media with glycerol, since PdeL levels are low in LB, but high in minimal media with an alternative carbon source such as glycerol (48). When strains were grown in either M9 or M63 minimal media supplemented with glycerol, we observed a ∼2-fold increase in P*_wec_* activity in the Δ*pdeL* strain compared to the wild type strain (**Fig. 1**). These data show that the P*_wec_* reporter can be used to detect changes in transcriptional regulation of the *wec* promoters. We therefore proceeded to test potential transcriptional regulators of the *wec* operon using the pP*_wec_*::*lux* reporter.

**Fig 1.**
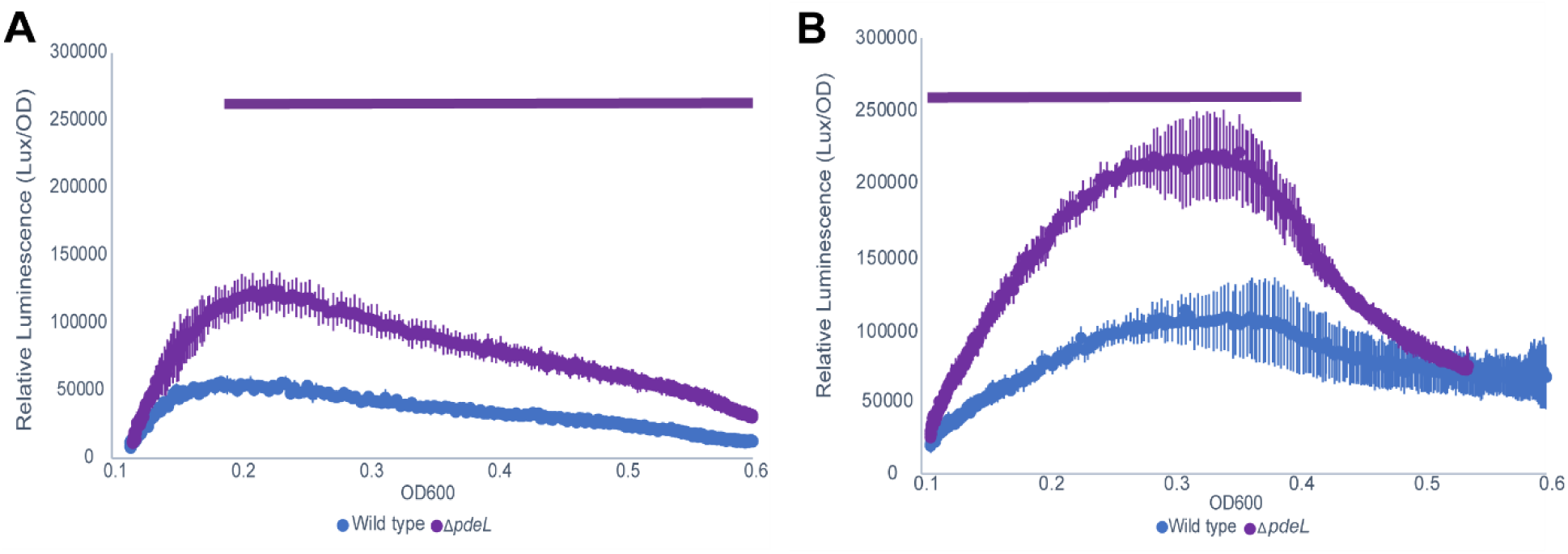
Reporter detects changes in P*_wec_* activity from a known transcriptional regulator. P*_wec_* promoter activity was measured by relative luminescence from the pJW15-P*_wec_* reporter plasmid in Δ*pdeL* strain. PdeL directly represses transcription of the *wec* operon (44). **A.** P*_wec_* activity was significantly increased in a Δ*pdeL* mutant compared to wild type when cells were grown in M9 glycerol minimal media. **B.** P*_wec_* activity in a Δ*pdeL* mutant was also increased compared to wild type when cells were grown in M63 glycerol minimal media, although reporter activity was higher in this media condition overall. Data are shown as luminescence/OD_600_ vs. OD_600_ to indicate growth phase for the mean of 3 biological replicates ± the SEM. Bar: Continuous time points where p<0.05 by Student’s T-test.

To identify candidate transcriptional regulators of the *wec* operon, we used PRODORIC’s Virtual Footprint tool (49) to cross-reference the *wec* promoter region (**Fig. S1**) (from 500 base pairs upstream of the start codon of the first gene in the operon (*wecA*) to 20 base pairs into *wecA*) against a library of *E. coli* binding site consensus sequence matrices for well-characterized *E. coli* transcriptional regulators (49, 50). Candidate regulators with moderate (0.70-0.85) and high relative similarity (≥ 0.85) scores were selected to be tested (**Tables S1 and S2**). To determine P*_wec_* activity in strains lacking the potential regulators, we constructed strains with single deletions of each of the candidate regulators (**Table S3**) and transformed them with the pP*_wec_*::*lux* reporter.

Of the 36 potential regulators identified, four could not be assayed due to non-specific genetic interactions with the reporter plasmid (*fis* and *oxyR*), severe growth defects in the mutant strains (*glnG*, *hns*), or activity in anaerobic conditions incompatible with the luciferase reporter (*arcA*). Of the remaining 32 potential regulators assayed, four mutants, Δ*gadX*, Δ*metJ*, Δ*nagC*, and Δ*phoB*, demonstrated statistically significant differences in P*_wec_* activity from the wild type strain (**Fig. 2A-E**). GadX is a transcriptional regulator of *gadE*, the master regulator of the glutamate-dependent acid resistance system (51). We observed increases in P*_wec_* activity in the Δ*gadX* both when cells were grown at a neutral pH of 7.0 (**Fig. 2A**) and at a mildly acidic pH 5.0 (**Fig. 2B**), although the effect of GadX was larger at pH 5 (approximately 2-fold). These data suggest GadX may be a repressor of P*_wec_*, especially at acidic pH’s. MetJ is a transcriptional repressor of methionine biosynthesis and is important for maintaining cellular redox balance (52). When methionine is abundant, a corepressor S-adenosylmethionine (SAM) binds to MetJ, which then represses the various methionine biosynthesis genes of the *met* regulon (52). In LB, the average concentration of L-methionine is high (2.0 mM) (53), and when we assayed Δ*metJ*, we found a 4-fold increase in *wec* promoter activity compared to the control, suggesting MetJ may be a repressor of *wec* activity (**Fig. 2C**). Deletion of *nagC*, which codes for a transcriptional regulator of the GlcNAc (*nag*) operon, resulted in a 2-fold increase in *wec* transcriptional activity (**Fig. 2D**). PhoB is the response regulator of the PhoRB two-component system which responds to phosphate limitation (54, 55). When grown in phosphate-limited conditions, a Δ*phoB* mutant demonstrated a 3-fold decrease in transcriptional activity compared to the wild type (**Fig. 2E**). We largely observed similar growth rates in the mutant strains compared to the wild type, excepting a small, but statistically significant, increase in growth in the Δ*phoB* mutant grown in MOPS minimal media and the Δ*gadX* mutant grown in LB at pH 7 (**Fig. S2**). The other 32 regulators tested did not significantly alter P*_wec_* activity compared to wild type in the conditions tested (**Fig. S3**); however, some might have an effect in other environmental conditions that were not tested. Nevertheless, we proceed by further analyzing the role of GadX, MetJ, NagC, and PhoB in regulation of the *wec* operon and ECA synthesis.

**Fig 2.**
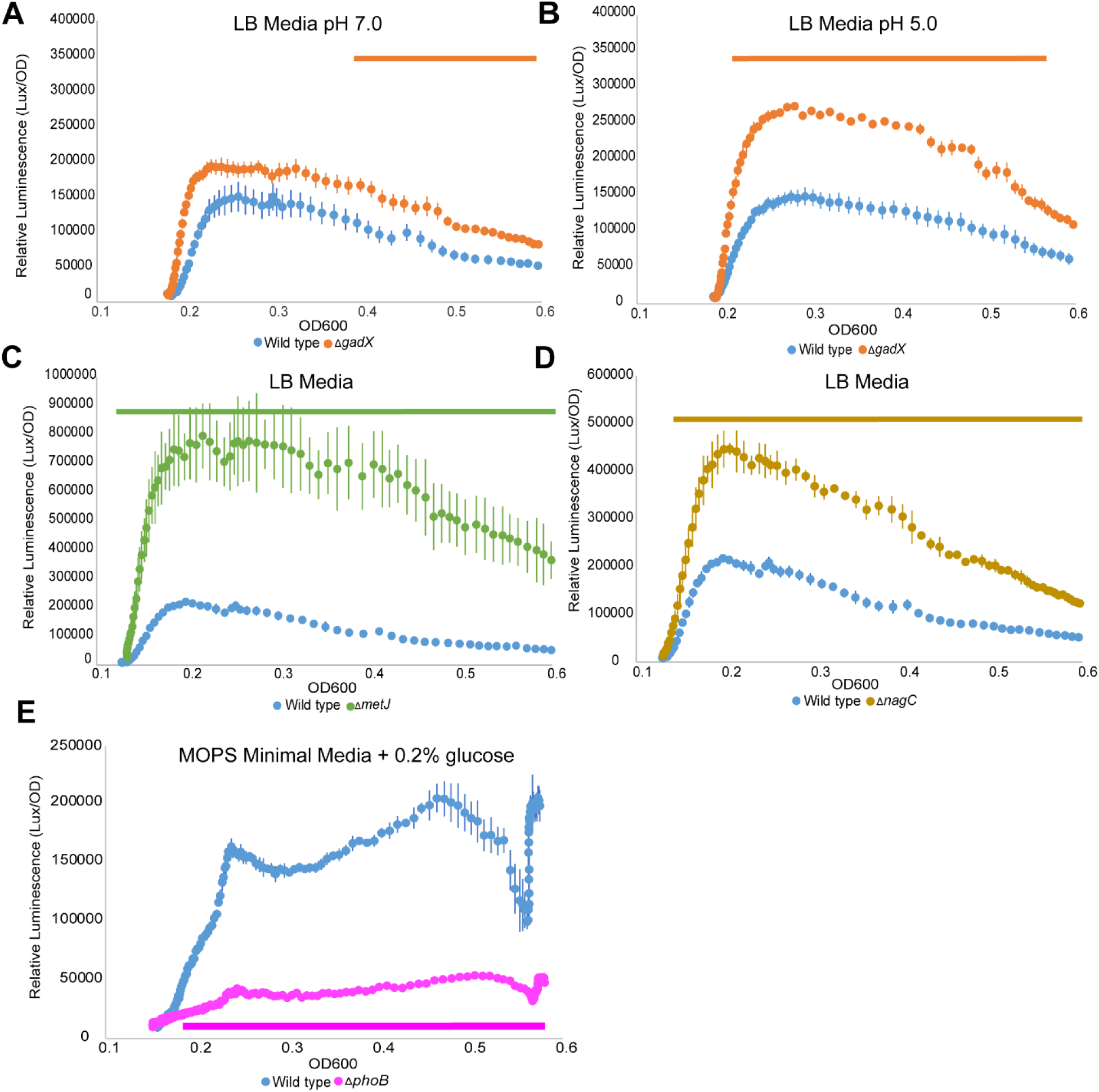
Loss of predicted transcriptional regulators alters P*_wec_* activity. For transcriptional reporter assays, luminescence and OD_600_ were measured over a six-hour period, except for minimal-media conditions which were run for 10 hours. **A.** The Δ*gadX* mutant increased P*_wec_* activity compared to the wild type in pH 7 non-buffered LB media. **B.** A larger increase in P*_wec_* activity was observed in a Δ*gadX* mutant in pH 5 non-buffered LB media. **C.** The Δ*metJ* mutant caused an increase in P*_wec_* activity compared to the wild type in LB media. **D.** The Δ*nagC* mutant had increased P*_wec_* activity in LB media. **E.** The Δ*phoB* mutant showed decreased P*_wec_* activity in low phosphate conditions (MOPS minimal media with 0.2% glucose). Corresponding growth curves for the wild-type and mutant strain are shown in **Fig. S2**. Data shown as luminescence/OD_600_ vs. OD_600_ for the mean of 3 biological replicates ± the SEM. Note, y-axis range differs between panels. Bar: Continuous time points where p<0.05 by Student’s T-test.

### MetJ and GadX directly bind P*_wec_ in vitro*

Although GadX, MetJ, NagC, and PhoB were identified based on potential binding sites within the P*_wec_* region, the change in P*_wec_* activity we observed may be due to direct regulation by the identified regulators or indirect regulation due to other transcriptional changes. To determine whether the transcriptional regulators could modulate P*_wec_* activity via direct binding, we conducted a series of electrophoretic mobility shift assay (EMSAs) where purified His-tagged protein was incubated with a 520-base pair P*_wec_* PCR product containing the primary and secondary *wec* transcriptional start sites (TSS) (see **Fig. S1**) (56). Virtual Footprint suggests three potential GadX binding sites in the P*_wec_* region, two of which overlap with the TATA boxes of the P*_wec_* promoters (**Fig. 3A**), consistent with a potential role as a repressor. For the *gadBC* promoter positive control, we selected a 224 base pair region upstream of *gadBC*, which contains three experimentally confirmed binding sites for GadX (57) (**Fig. 3B**). When we assayed GadX binding to the P*_wec_* promoter region, we observed a mobility shift consistent with binding of GadX to the P*_wec_* DNA (lane 1 vs. 3) (**Fig. 3C**) and similarly observed a mobility shift for our DNA positive control (**Fig. 3C**, lane 4 vs. 5). GadX interaction with P*_wec_* was maintained even in the presence of a non-specific competitor poly(dI-dC) (**Fig. 3C**, lane 10) or the specific competitor positive control DNA (**Fig. 3C**, lane 6), suggesting *in vitro* GadX binding at P*_wec_* is specific and its affinity is at least as strong as for the *gadBC* promoter.

**Fig. 3.**
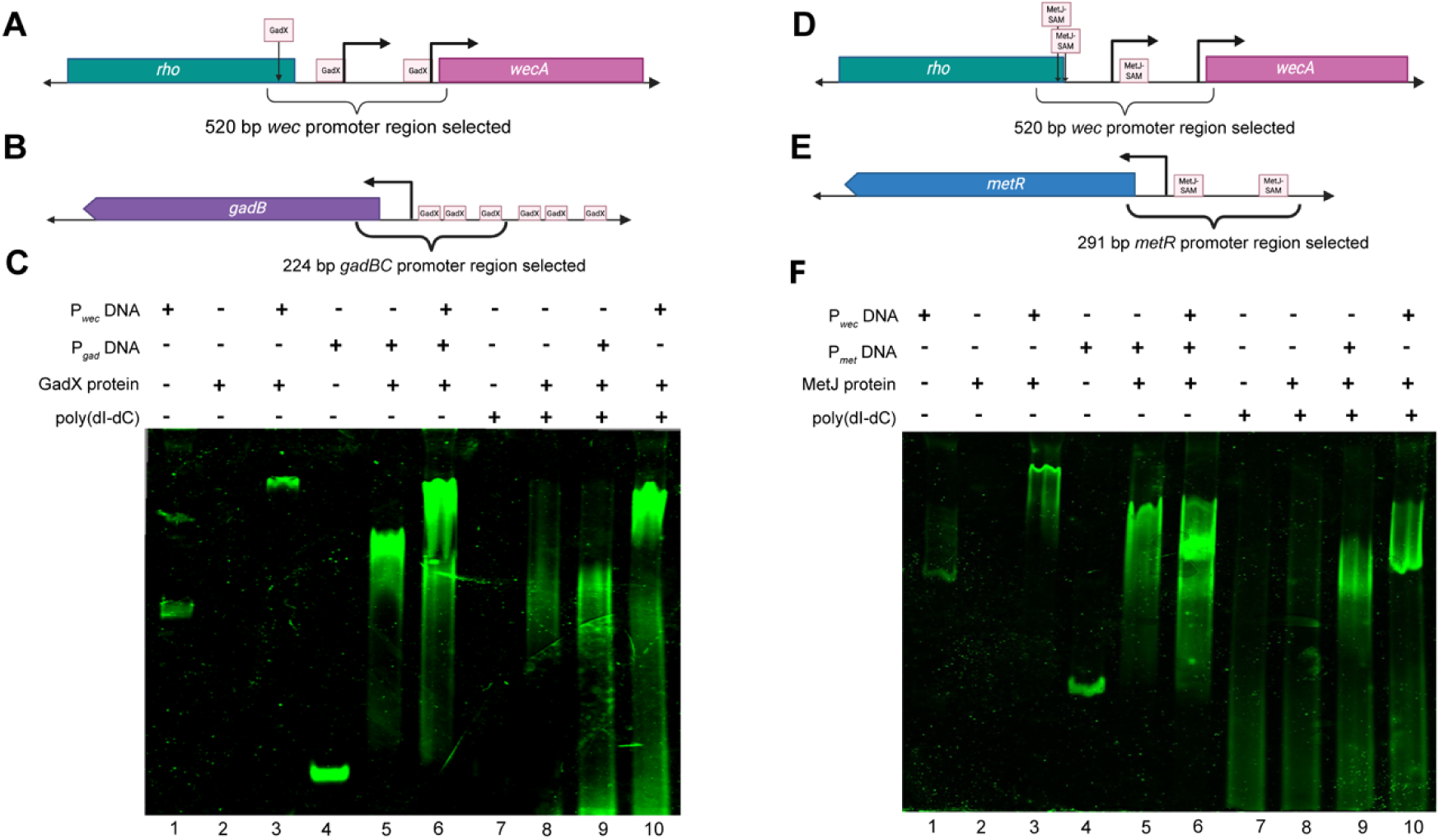
GadX and MetJ bind the *wec* operon promoter region *in vitro*. **A.** Graphic showing the three predicted GadX binding sites in the *wec* promoter region from PRODORIC Virtual Footprint (49). **B.** Graphic showing known GadX binding sites in the positive control DNA, a 224-base pair region upstream of the *gadBC* operon (P*_gad_*). **C.** EMSA conducted with 100 nM of purified His-GadX per lane and 15 nM of target DNA per lane where indicated. GadX binds to P*_wec_* DNA causing a size shift (Lane 3 vs. 1) and this shift is maintained both in the presence of specific (P*_gad_*) and non-specific competitor (poly(dI-dC)) DNAs (Lane 6 and 10, respectively). **D.** Graphic showing the three predicted MetJ binding sites in the *wec* promoter region from PRODORIC Virtual Footprint. **E.** Graphic showing known MetJ binding sites in the positive control DNA, a 291-base pair region upstream of *metR* (P*_met_*). **F.** EMSA was performed as in (C), except that SAM was added to potentiate MetJ DNA binding. MetJ binds to P*_wec_* DNA causing a size shift (Lane 3 vs. 1). A partial shift is maintained in the presence of specific (P*_met_*, Lane 6) or non-specific (poly(dI-dC), Lane 10) competitor DNAs, suggesting that MetJ binding to a subset of binding site(s) on the P*_wec_* DNA is specific and robust. A similar partial shift is observed for the positive control DNA (Lane 9). Images are representative of 3 independent experiments. Panels A, B, D, and E created in BioRender. Hill, H. (2026) https://BioRender.com/3k9j47h

We then tested the ability of MetJ to bind to the P*_wec_* region. SAM was added to the MetJ binding reactions, since SAM acts as a transcriptional corepressor, enhancing binding at *met* boxes by 10-fold (58). MetJ was also predicted to have three potential binding sites in the P*_wec_* region, which are located close to the predicted transcriptional start sites (**Fig. 3D**). For the *met* promoter positive control, we selected a 291 base pair region upstream of *metR*, which contains two experimentally confirmed binding sites for MetJ-SAM (**Fig. 3E**) (57). We observed a mobility shift when from the P*_wec_* DNA was incubated with MetJ-SAM (**Fig. 3F**, lane 1 vs. 3), suggesting binding of MetJ-SAM to the P*_wec_* region. We observed a similar mobility shift when the positive control DNA was incubated with MetJ-SAM (**Fig. 3F**, lanes 4 vs. 5). Interestingly, when we incubated the P*_wec_* DNA with MetJ-SAM in the presence of the specific competitor (lane 6) or non-specific competitor (lane 10), we observed a smaller mobility shift than that without a competitor (lane 3). This suggests that binding to some but not all binding sites on the P*_wec_* DNA is specific. We observed a similar lesser shift in the band size of the positive control DNA (lanes 6 and 9 vs. 5), suggesting the same is true for the *met* promoter region. As the P*_wec_* DNA behaves similarly to the positive control DNA, our results suggest that MetJ-SAM directly interacts with P*_wec_ in vitro*. An EMSA conducted for NagC did not show significant binding of NagC to the P*_wec_* DNA (**Fig. S4**), suggesting regulation of P*_wec_* activity by NagC may be indirect. We were not successful in purifying functional PhoB protein and so could not conduct an EMSA for PhoB binding to P*_wec_*. Overall, these data demonstrate that GadX and MetJ can bind to P*_wec_ in vitro* and suggest a model where the altered P*_wec_* activity we observed in the Δ*gadX* and Δ*metJ* mutants is due to loss of direct binding by these proteins at operator sites in P*_wec_*.

### Changes in P*_wec_* activity with Δ*gadX* and Δ*metJ* are reflected in linear ECA accumulation

ECA immunoblots were conducted to determine if the level and direction of changes in P*_wec_* activity observed in the transcriptional reporter data correlated with steady-state levels of linear ECA synthesized (ECA_CYC_ cannot be resolved by immunoblotting (40)). The levels of linear ECA are much higher in stationary phase than during exponential growth (**Fig. S5A**), possibly due to the large effect of RpoS on ECA levels (**Fig. 4AB**, **Fig. S5BC**). Therefore, we conducted immunoblots from mid-log phase (**Fig. 4AB**) and stationary phase (**Fig. S5BC**) samples. Both the Δ*metJ* and Δ*gadX* strains had increased P*_wec_* activity compared to the wild type (**Fig. 2A-C**).

**Fig. 4.**
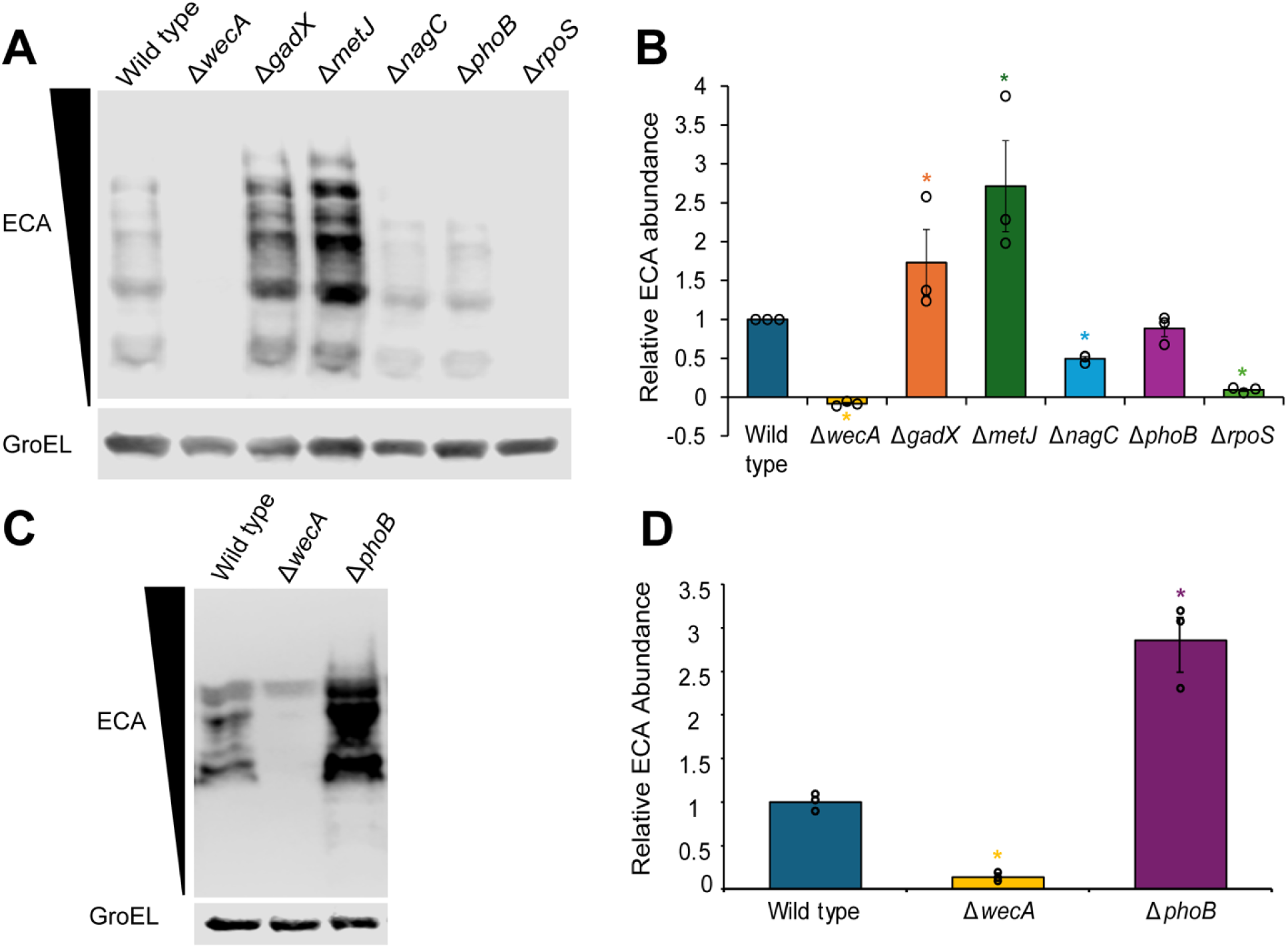
Regulation deletion alters ECA abundance. **A.** ECA immunoblot conducted using mid-log phase (0.5 OD_600_) wild-type and regulator gene deletion strains grown in LB media. GroEL acts as normalization control, while Δ*wecA* is a negative control for ECA (showing any non-specific bands). **B.** Quantification of ECA immunoblot in (A) normalized to wild type ECA levels. All regulator mutants show significant changes in ECA accumulation except Δ*phoB*. **C.** ECA immunoblot conducted using stationary phase cultures grown in MOPS with 0.2% glucose as a low phosphate condition. GroEL acts as a normalization control, while Δ*wecA* serves as a negative control. **D.** Quantification of ECA immunoblot in (C). The Δ*phoB* mutant has a significantly higher level of ECA than the wild type, in contrast to the transcriptional regulation. Images are representative of three independent experiments. Quantitative data are shown as the mean of 3 biological replicates ± the SEM and individual data points. * p<0.05 by Wilcoxon test with normal approximation

Consistent with this result, we observed increased accumulation of ECA in mid-log samples from these strains (**Fig. 4**). In stationary phase, the change in ECA levels was smaller and significant only for Δ*metJ* and not for Δ*gadX* (**Fig. S5BC**). This may suggest that the effect of RpoS on ECA levels in stationary phase overwhelms that of the other regulators. While Δ*nagC* caused increased P*_wec_* activity, ECA levels were significantly decreased in the Δ*nagC* mutant compared to wild type in both log phase and stationary phase (**Fig. 4AB**, **Fig. S5BC**), although the change was smaller in stationary phase.

NagC represses genes necessary for the catabolism of GlcNAc (59). Therefore, decreased cellular abundance of GlcNAc may inhibit ECA synthesis in the Δ*nagC* strain even in the presence of increased P*_wec_* activity. As phosphate levels in LB are well above those that deactivate the PhoRB system (60–62), we did not observe a significant change in ECA levels with Δ*phoB* in either log phase or stationary phase samples grown in LB (**Fig. 4AB**, **Fig. S5BC**). Therefore, we grew wild-type and Δ*phoB* cultures to mid-log and stationary phase in low phosphate conditions (MOPS media (61, 62)) and assayed ECA levels. Contrary to our expectations, we found the Δ*phoB* strain had higher ECA levels at stationary phase than did the wild-type strain (**Fig. 4CD**). However, in actively growing cells, very little linear ECA was detectable in all tested strains in low phosphate conditions. It is unclear why the levels of ECA do not reflect the regulation of P*_wec_* activity for PhoB. Overall, these data demonstrate that the regulators that can bind P*_wec_* directly affect ECA levels in a manner consistent with their transcriptional effects, whereas the effects of the other regulators are more complex.

### ECA supports efficient growth in acidic conditions

We next proceeded to investigate genetic interactions between ECA and Δ*gadX* and Δ*metJ,* the regulators that could bind P*_wec_* and had consistent transcriptional regulation and ECA levels. We first asked whether GadX and MetJ affected ECA’s role in the OM permeability barrier. We tested single and double mutants lacking the GadX or MetJ and all ECA (Δ*wecA*) or only ECA_CYC_ (Δ*wzzE*) for resistance to compounds for which ECA impedes their cell entry, including bile salts, deoxycholate (a bile salt derivative), and SDS-EDTA (16). However, we did not observe any shifts in resistance indicating genetic interactions (**Fig. S6**). These results suggested that GadX and MetJ do not link ECA to bulk envelope permeability and raised the possibility that these regulators couple ECA production to the specific stresses under which they are active. We therefore turned our investigations to whether ECA plays a role in response to acidic conditions (GadX) and RNS stress (MetJ). We conducted transcriptional reporter assays on wild-type cells grown in unbuffered LB media adjusted to varying pH and observed that as pH decreases, the P*_wec_* activity increases **(Fig. 5A)**. This change in P*_wec_* activity did not, however, occur when cells were grown overnight at the assay pH before measuring P*_wec_* activity, habituating them to growth in the acid pH (**Fig. 5B**). These data indicate that a downshift in pH leads to induction of P*_wec_* activity but that this increased activity is not maintained over long time scales.

**Fig. 5.**
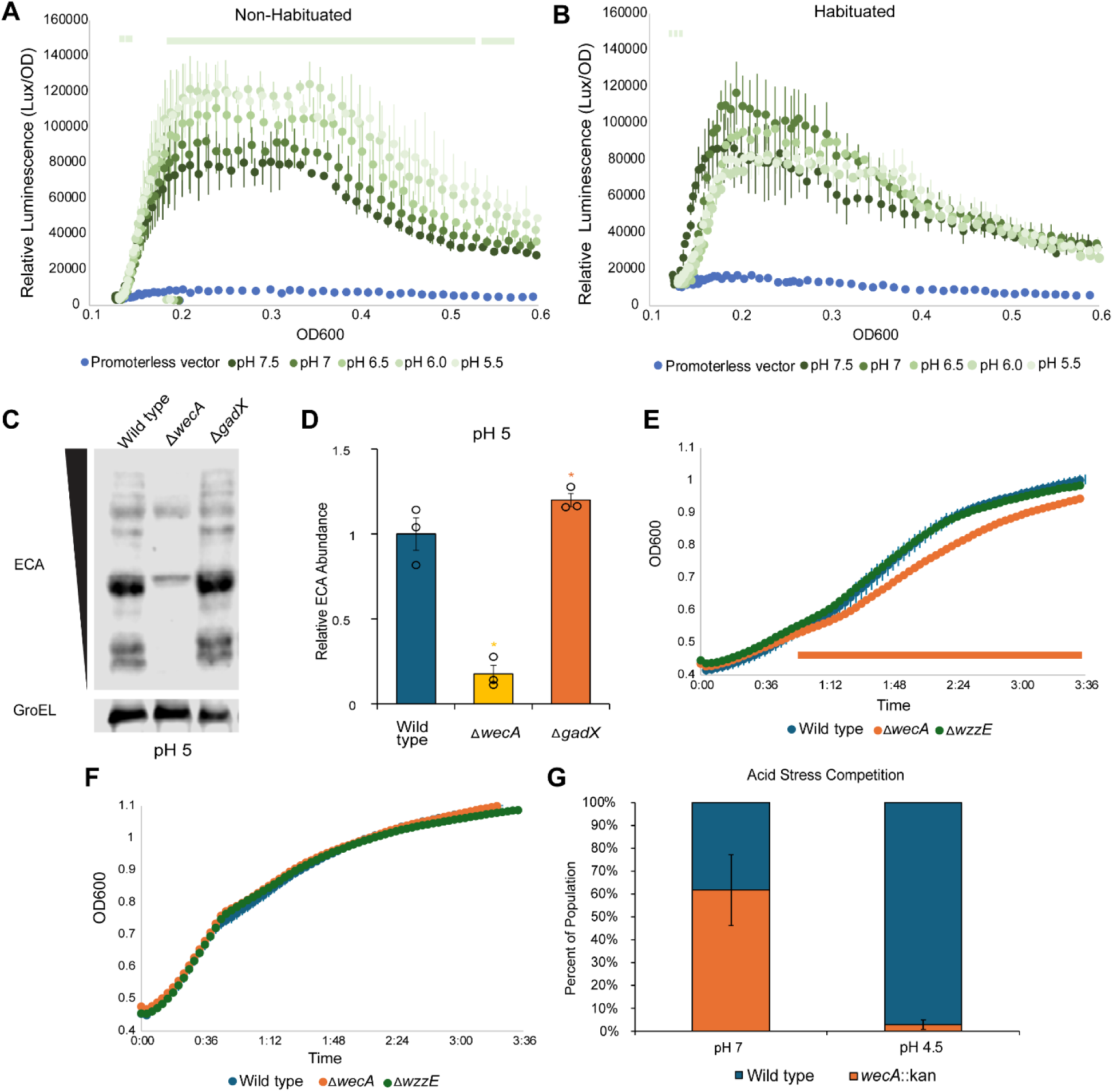
ECA plays a role in supporting growth at acidic pH. **A.** A P*_wec_* transcriptional reporter assay of non-habituated wild-type cells (grown overnight at neutral pH before back-dilution into media at the indicated pHs) shows that P*_wec_* activity increases at lower pHs in these conditions. **B.** In contrast, there is no significant difference in P*_wec_* activity when cells are habituated in the assay pH overnight before the transcriptional reporter assay. Data are shown as the mean of 3 biological replicates ± the SEM. Bar: p<0.05 of pH 5.5 vs. pH 7.5 by Student’s T-test. **C.** ECA immunoblot comparing wild-type and Δ*gadX* strains grown to 0.5 OD_600_ in mildly acidic pH (5.0). Δ*wecA* serves as a negative control for ECA (showing non-specific bands), while GroEL serves as a normalization control. **D.** Quantification of ECA immunoblot normalized to wild-type ECA levels. The Δ*gadX* strain shows a small, but significant, increase in ECA levels over those of wild type, but less so than at pH 7. Data are shown as the mean of 3 biological replicates ± the SEM and individual data points. * p<0.05 by Wilcoxon test with normal approximation. **E.** Growth curves were conducted at pH 4.5. The Δ*wecA* strain, which lacks ECA, shows slower growth at this pH than does the wild-type strain; however, the Δ*wzzE* strain (no ECA_CYC_, unregulated linear chain length) grows comparably to wild type at pH 4.5. **F.** The Δ*wecA* strain does not have a growth defect when grown at pH 7.0. Data are shown as the mean of 3 biological replicates ± the SEM. Bar: p<0.05 for Δ*wecA* vs. wild type by Student’s T-test. **G.** In a 1:1 competition assay, *wecA*::kan cells showed a significant fitness defect compared to wild type cells at pH 4.5 but not at pH 7. Data are shown as the mean of 3 biological replicates ± the SEM. * p<0.05 for *wecA*::kan vs. wild type in pH 4.5 by Student’s T-test.

Given the differing effects of pH on P*_wec_* activity at short and long-time scales, we then asked what effect GadX has on ECA levels at acidic pH. Therefore, we grew cells in acidic conditions (pH 5.0) and assayed the effect of Δ*gadX* on ECA levels.

Interestingly, we observed less change in ECA levels in the *ΔgadX* mutant compared to wild type at an acidic pH (**Fig. 5CD**) than we did at a neutral pH (**Fig. 4**). These data suggest that the repression of P*_wec_* activity by GadX is relieved in this case—perhaps by the activity of another acid stress regulator. When we assayed the effect of two other acid stress regulators (51) on P*_wec_* activity under acidic conditions, we found that Δ*gadW* did not change P*_wec_* activity, but increasing RpoS levels by removing a degradation adaptor (*sprE*::Tn*10* (63)) increased P*_wec_* activity and deleting *rpoS* in this background lead to very low levels of P*_wec_* activity (**Fig. S7)**. Given that Δ*rpoS* also had a large effect on ECA levels (**Fig. 4**, **S5**), these data suggest that RpoS may be responsible for the increase in P*_wec_* activity in acidic pH. GadX effects on ECA levels are most readily detected at neutral pH, suggesting direct repression by GadX fine tunes *wec* expression rather than driving its induction in acidic conditions.

The increase in P*_wec_* activity at acidic pH suggests that ECA may play a role in helping cells adapt to growth in acidic conditions. Therefore, we performed growth curves where wild-type cells or cells lacking all ECA (Δ*wecA*) or only ECA_CYC_ (Δ*wzzE*) were grown to mid-log phase in neutral conditions then transferred to acidic conditions to assay growth and survival. We observed that growth at pH 4.5 was attenuated in the Δ*wecA* mutant compared with wild type cells (**Fig. 5E**), whereas all strains grew well at neutral pH (**Fig. 5F**). To determine whether the Δ*wecA* growth defect resulted in decreased competitive fitness, we conducted a competition assay between ECA-lacking cells (Δ*wecA*::kan) and wild type cells with equivalent starting CFUs/mL (**Fig. 5G**). We observed that in cells grown at neutral pH and plated at mid-log, Δ*wecA*::kan cells and wild type cells did not show a significant difference in fitness. However, during acid stress (pH 4.5), the ECA-lacking cells were significantly outcompeted by the wild type. Together, these data indicate that a linear form(s) of ECA is important for fitness in acidic conditions.

### ECA is required for optimal growth under RNS stress

GSNO is a source of bioavailable nitric oxide produced by hosts during infection that induces RNS and causes protein S-nitrosylation (64). In *E. coli*, GSNO nitrosates the thiol groups of intracellular homocysteine and cysteine, draining intermediates in the methionine biosynthesis pathway (65). Thus, MetJ is a central regulator of the GSNO response: GSNO exposure alters MetJ regulatory activity, causing a large increase in expression of *met* genes (e.g., *metE*, *metF*, *metB*) (65). Microarray data suggests *wecA (rfe)* expression may change in response to GSNO in a MetJ-dependent manner (65).

Therefore, we aimed to determine the effect of GSNO on P*_wec_* activity in wild-type and Δ*metJ* cells. As in **Fig. 2C**, in vehicle-treated cells, the P*_wec_* activity in a Δ*metJ* strain is higher than that of the wild-type strain (**Fig. 6A**). When treated with 100 μM GSNO, the wild-type P*_wec_* activity increases to a level similar to that of the Δ*metJ* strain with or without treatment (**Fig. 6B**). This suggests that GSNO relieves repression by MetJ in the wild-type strain, causing increased *wec* expression.

**Fig. 6.**
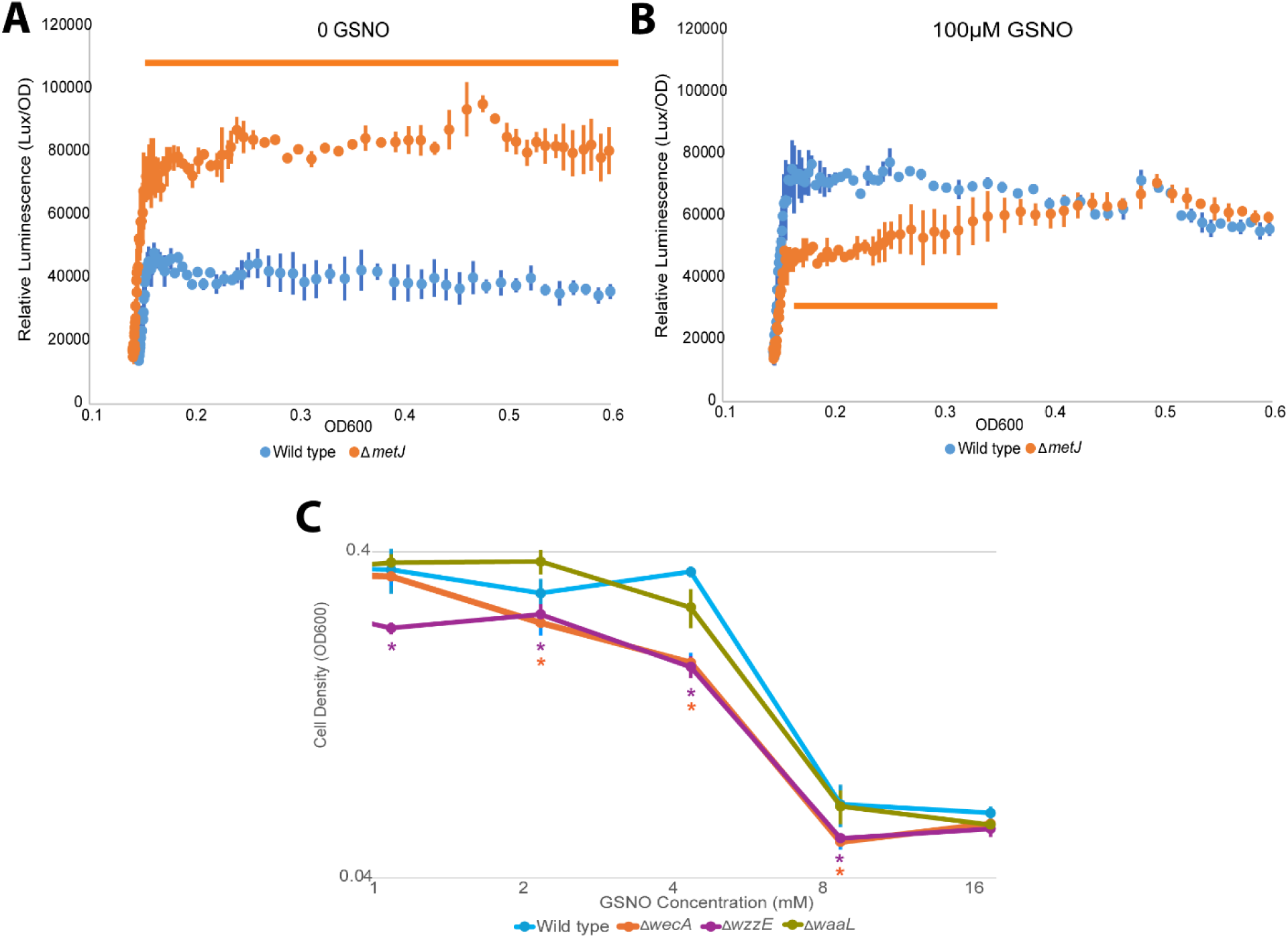
ECA plays a role in growth under RNS stress. GSNO is a bioavailable form of nitric oxide and causes RNS stress. **A.** P*_wec_* activity in media without GSNO is significantly higher in the Δ*metJ* strain than in wild type. **B.** Under RNS stress (100 μM GSNO), P*_wec_* activity in the wild type strain increases to phenocopy that of the Δ*metJ* strain. Bar: p<0.05 by Student’s T-test. **C.** Cells were grown for 24 hours at 37 °C with the indicated concentration of GSNO. Higher GSNO concentrations are used to induce growth inhibition. Loss of all ECA (Δ*wecA*) or of ECA_CYC_ and chain-length regulation (Δ*wzzE*) resulted in less growth under RNS stress. * p<0.05 for Δ*wecA* vs. wild type (blue) or Δ*wzzE* vs. wild type (green). Statistical significance determined by one-way ANOVA with Dunnett’s post-hoc test. All experiments were performed in M63 glucose media. Data are shown as the mean of 3 biological replicates ± the SEM.

Interestingly, the *wec* operon is predicted to be regulated by another RNS stress regulator, NsrR (66, 67). When we tested P*_wec_* activity in a Δ*nsrR* strain, we observed a non-statistically significant, but repeatable, increase in P*_wec_* activity compared to the wild-type strain (**Fig. S8A**). When we treated with 100 μM GSNO, wild-type P*_wec_* activity increased to the level of the Δ*nsrR* strain, while P*_wec_* activity in this strain did not change (**Fig. S8B**). Although the increase in P*_wec_* activity in the Δ*nsrR* did not reach statistical significance, it was reproducible across replicates and its response paralleled that of the Δ*metJ* strain. Therefore, these data, combined with the Δ*metJ* data suggest that ECA may play a role in resistance to RNS stress.

Therefore, we sought to determine whether ECA would affect growth or survival under GSNO treatment. Accordingly, we grew wild-type, Δ*wecA*, and Δ*wzzE* cells with various concentrations of GSNO and measured their final density after 24 hours. Both the Δ*wecA* (no ECA) and Δ*wzzE* (no ECA_CYC_) mutants had a statistically significant reduction in growth at concentrations of GSNO compared to the wild-type in an MIC-style assay (**Fig. 6C**), while the Δ*waaL* (no ECA_LPS_) strain behaved identically to the wild-type strain (**Fig. 6C**). Because the mutant lacking ECA_LPS_ grew similarly to wild type whereas both mutants lacking all ECA and lacking ECA_CYC_ were impaired, the RNS growth defect is specific to loss of ECA_CYC_ or the regulation of linear ECA chain length rather than ECA synthesis in general. Overall, our data demonstrate ECA contributes measurably to growth under the environmental stresses that activate its transcriptional regulators, linking ECA production to acid and RNS stress responses.

## Discussion

This study identified GadX and MetJ as repressors of the *E. coli wec* operon capable of binding P*_wec_ in vitro* and implicates ECA in adaptation to acid and RNS stress. Prior to this work, the only experimentally confirmed transcriptional regulator of the *wec* operon was PdeL, a cyclic di-GMP phosphodiesterase and transcriptional regulator (44, 48). Therefore, our work substantially broadens the known regulatory landscape of ECA biosynthesis and reveals surprising functions for the GadX and MetJ regulators that have been characterized in processes unrelated to glycan metabolism. Moreover, these findings provide evidence that the synthesis of a highly conserved *Enterobacterales* glycan is dynamically regulated, tuning envelope remodeling to specific environmental stresses.

GadX canonically functions within a complex signal transduction network governing the glutamate-dependent acid resistance response (see **Fig. S9**). GadX and GadW are AraC-family regulators that induce expression of glutamate decarboxylase genes via transcriptional regulation of the central regulator, *gadE* (68). In enteropathogenic *E. coli*, GadX also participates in transcriptional control of plasmid-encoded virulence genes (69), linking pH sensing to coordinated control of acid resistance and virulence factor expression. In this network, PhoP activates transcription of *iraM*, whose product binds SprE, preventing SprE from targeting RpoS to the ClpXP protease for degradation, thereby allowing RpoS to activate *gadE* transcription (70). RpoS then activates GadY, a small RNA that stabilizes GadX mRNA (71).

Our data show that GadX represses P*_wec_* activity and that the effect of this repression on ECA abundance is less at lower pHs, despite the activation of GadX. This supports a model in which another regulator is responsible for the increased P*_wec_* activity we observe initially at lower pH and then GadX fine-tunes *wec* expression (**Fig. S9**). We propose that, under neutral conditions, GadX contributes to baseline repression of the *wec* operon. Upon acidification, RpoS levels increase through the actions of IraM and SprE and cause increased *wec* transcription, while GadX-dependent repression follows, partially counterbalancing this induction and moderating ECA levels. This model provides a potential explanation for the loss of P*_wec_* induction in acidic pH after habituation: continued GadX activation may return P*_wec_* activity to baseline. Future analyses of *wec* transcription over time in acidic conditions, and of regulator interactions, could expand on the mechanisms that lead to the change in P*_wec_* activity over time.

Nevertheless, the large fitness defect observed for ECA-deficient cells in acidic conditions suggests that ECA is regulated under acidic conditions in order to promote the cell’s fitness. That we observed this fitness defect in a Δ*wecA* strain but not a Δ*wzzE* strain suggests that the linear forms of ECA are involved in maintaining growth in acid conditions. Interestingly, in a pathogenic Shiga-toxin producing *E. coli* strain (O157:H7), mutants lacking ECA and O-antigen have been shown to be defective for growth in concentrations of acetic acid that produce mildly acidic pH, while a mutant lacking only ECA displayed partial resistance (72). These data suggest that ECA’s integration into the acid stress response extends beyond lab strain *E. coli* and may in fact be more pronounced in some pathogenic strains or by the acid source (i.e., organic acids). The role of ECA in promoting growth in different acid sources and in clinically relevant strains are interesting avenues for future studies.

Our results also extend prior transcriptomic evidence (65) connecting GSNO exposure to altered *wecA* expression by identifying MetJ as a repressor of the *wec* operon and by demonstrating that ECA biosynthetic mutants Δ*wecA* and Δ*wzzE* have reduced growth during RNS stress. MetJ is best known as a SAM-dependent regulator of the methionine-biosynthesis (*met*) regulon (73, 74). MetJ and the other primary *met* regulator, MetR, jointly control expression of the *met* regulon (57, 75). Beyond methionine biosynthesis, this integrated MetJ/MetR system is incorporated into broader environmental feedback pathways, including a role in RNS protection (52, 65, 76). Methionine biosynthesis contributes to cellular redox balance by acting as a “redox sink” detoxifying oxidants (77, 78); however, RNS, such as GSNO, inhibit the function of the methionine synthases (MetE and MetH) lowering methionine levels and disrupting redox balance (65, 79, 80). In this context, RNS decrease methionine and thus SAM levels derepressing MetJ-regulated genes.

We observed this derepression for P*_wec_* activity with GSNO treatment, even at levels far lower than those required to cause growth inhibition, further supporting the integration of ECA into the RNS stress response. As further suggestive evidence of ECA’s role in resisting RNS stress, we also observe a repeatable effect of NsrR on P*_wec_* activity that is relieved by GSNO treatment. Overall, our findings support a model in which MetJ directly represses *wec* operon transcription, linking methionine and SAM homeostasis to ECA production (**Fig. S10**). We propose that, under conditions which support MetJ-SAM-dependent repression, MetJ binds within the P*_wec_* region and restricts *wec* expression. During GSNO exposure, MetJ-dependent repression is inhibited and P*_wec_* activity is increased (**Fig. S10**). Increased ECA abundance then helps to preserve growth under higher, growth inhibiting concentrations of RNS. Interestingly, *ispU*, the gene responsible for synthesizing Und-P (81), which is used for synthesis of extracytoplasmic glycans including ECA, has also been suggested to be regulated in response to GSNO treatment (65), suggesting a larger connection between RNS and extracytoplasmic glycans. Unlike for acid stress, loss of either all ECA or loss of ECA_CYC_ and linear chain length regulation caused a growth defect under RNS stress. The changes in ECA levels caused by regulator deletion were not sufficient to alter OM permeability phenotypes. Together, these data demonstrate that different properties of the ECA glycan are necessary for different cellular functions, a finding that could expand understanding of carbohydrate structure-function relationships in the future. The mechanism behind ECA’s effect on growth in RNS and acid stress will be an intriguing area for future investigation.

In contrast to GadX and MetJ, our NagC and PhoB results demonstrate that P*_wec_* activity does not always predict accumulated linear ECA, as deletion of the genes for both regulators resulted in opposing changes in P*_wec_* activity and ECA levels. NagC represses genes involved in the catabolism of GlcNAc (59, 82–84), suggesting that the lower ECA levels observed in a Δ*nagC* strain are due to low cellular availability of GlcNAc. Further, as NagC did not bind *in vitro* to P*wec*, one possible explanation is that the increased P*_wec_* activity in a Δ*nagC* strain is due to compensatory regulation in response to the low ECA levels. As we could not test PhoB binding to P*_wec_*, the mechanism underlying the opposing phenotypes of the Δ*phoB* strain is less clear. However, it may be that this effect is also mediated by substrate availability or other biosynthetic constraints. Together, these findings indicate that ECA accumulation is influenced by factors beyond *wec* transcription, suggesting that multiple layers of regulation interact to control ECA abundance.

In summary, our data expand the regulatory framework for ECA biosynthesis and show ECA contributes to fitness during acid and growth under RNS stress. The growth contributions of ECA we observed were most apparent during acute acid downshift and at intermediate concentrations of RNS, consistent with a role in fine-tuning envelope resilience rather than in acute survival. Nevertheless, as evidenced by the lack of competitive fitness of a mutant lacking ECA in acidic conditions, these effects may be highly significant during the chronic or competitive conditions *E. coli* encounters in their natural niches. GadX and MetJ are supported as repressors of *wec* transcription that can directly bind P*_wec_ in vitro*. Changing environmental conditions can damage the cell envelope through disruptions to membranes, protein folding, and/or biosynthetic pathways, and both envelope and non-envelope stress responses can remodel the cell envelope to counter those stresses (85). Our data integrate ECA into these global stress response networks and provide a new facet of cell envelope remodeling under stress. The detailed mechanisms of ECA regulation (including validation of the predicted regulator binding sites) and of its role in stress resistance will provide fascinating areas for future studies.

## Materials and Methods

### Bacterial strains and growth conditions

*E. coli* K-12 MG1655 was used in all experiments. Unless specified, overnight cultures were grown in LB Lennox medium for 16 hours at 37°C. When necessary for plasmid maintenance, the LB was supplemented with 25 μg/mL kanamycin (Gold Biotechnology) or 20 μg/mL chloramphenicol (Gold Biotechnology). All strains used in this study are listed in **Table S1**. Knockout strains in MG1655 were constructed with Keio collection alleles (86) using P1vir transduction (87). The pCP20 plasmid, which encodes a FLP recombinase, was used to remove resistance cassettes after transduction (88).

### Reporter assays

To quantify P*_wec_* activity, cultures of strains expressing the pP*_wec_*::*lux* reporter plasmid were grown overnight, and the next day strains were subcultured at 1:100 dilution and appropriate antibiotic was added for selection. The luciferase luminescence assay was performed as previously reported (45), with minor modifications. Specifically, for *metJ* GSNO transcriptional assays, a 48-well clear plate was used instead of a 24-well plate, with appropriate culture volume adjustment.

### Efficiency of plating assays (EOPs)

EOPs were performed as previously reported (16).

### ECA immunoblots

ECA immunoblots were performed as previously reported (16), with minor modifications. Specifically, chemiluminescence was imaged using a LI-COR Odyssey M imager and the results were quantified using the Image Studio version 6.0 software (LI-COR Biosciences).

### Protein expression and purification

For histidine-tagged protein isolation and purification, we expressed His-tagged genes of interest from an IPTG-inducible overexpression plasmid from the appropriate ASKA collection (89) strain using the following technique. 1 L volumes of LB media with chloramphenicol and 25 μM IPTG were inoculated with the respective strain and grown for 16 hours. Cultures were spun down at 5000 x g for 10 minutes and cells were resuspended in 5 mL BugBuster (Novagen) lysis buffer containing Benzonase (1:100) (Millipore Sigma) per 1 g wet cell paste. Cells were then incubated for 15 minutes at room temperature and an equal volume of equilibration buffer (50 mM NaH_2_PO_4_, 300 mM NaCl, 10 mM imidazole pH 8.0) was added and the solution was mixed. The lysate was spun down at 5000 x g for another 10 minutes. Then the protein-containing supernatant was filtered using 0.22-µm filter and kept at 4 °C prior to and during fast protein liquid chromatography (FPLC) on a BioRad NGC instrument. Protein solution was added to an EconoFit Nuvia IMAC Ni-charged column (BioRad) and His-tagged proteins were eluted using a gradient of elution buffer (50 mM NaH_2_PO_4_, 300 mM NaCl, 500 mM imidazole pH 8.0). The eluted protein solution was collected and ultrafiltered using Vivaspin 500 columns (Sartorius) to concentrate the protein and remove the imidazole. Proteins were stored in 20% glycerol at -80 °C.

### Electrophoretic mobility shift assay (EMSA)

100nM of purified protein of interest was incubated in a reaction mixture with 15 nM purified *wec* promoter DNA, 5 mM MgCl_2_, 0.5% NP-40, and a binding buffer containing 50 mM Tris-HCl (pH 7.5), 250 mM KCl, 5 mM EDTA, and 50% glycerol at room temperature for 20 minutes. In addition, 50μM of the corepressor S-adenosylmethionine (SAM) was added to the MetJ reaction to enhance binding. For the competition reactions, 250ng of the non-specific competitor poly(dIdC) was used. After incubation, loading dye was added and samples were electrophoresed in 0.5x TBE buffer on a 5% non-denaturing acrylamide gel (29:1 acrylamide:bis-acrylamide) for 90 minutes at 4°C. DNA was then stained with SYBR Green (Invitrogen) and imaged using a LI-COR Odyssey M at excitation 488 nm and filter settings appropriate for visualizing SYBR Green.

### Acid resistance assay

Overnight cultures of wild-type, Δ*wecA*, and Δ*wzzE* strains were sub-cultured in neutral pH LB media and grown to mid-log phase (0.5 OD_600_). Mid-log phase cells were then spun down for 5 minutes at 5,000xg and resuspended in the same volume of low pH LB medium (pH 4.5) and grown with shaking in a clear 48-well plate at 37°C in a BioTek Synergy H1 plate reader with OD_600_ measurements collected every 3 minutes.

### Acid Stress Competition Assay

Overnight cultures of wild-type and *wecA*::kan cells were normalized to an OD600 of 1.0, combined in a 1:1 ratio, and inoculated at a 1:100 dilution into 5mLs of either pH 7 liquid LB medium or pH 4.5 LB medium. The cultures were grown at 37°C until the culture reached an OD600 of 0.5; then, samples were serially diluted down to an expected concentration of 500 CFU/mL and plated on both plain LB and LB with 25 μg/mL kanamycin. Colonies were counted using an eCount™ Colony Counter Pen. Wild-type CFU was calculated by subtracting the kanamycin resistant count from the plain LB count.

## Acknowledgments

We thank the members of the Mitchell lab, Dr. Jolene Ramsey and her lab, Dr. Deborah Siegele, Dr. Beiyan Nan (Texas A&M University), and Dr. Anna Konovalova (UT Health Houston) for feedback, constructive criticism, and productive discussions of this project. We also thank Ms. Elizabeth Ponder for her contributions to strain building, Ms. Kavya Khandelwal for her experimental help, and Dr. Guadalupe Valencia-Toxqui for her assistance with the FPLC protein purification. Additionally, we thank Dr. Deborah Siegele for the wild type *E. coli* MG1655 K-12 strain, Dr. Tracy Raivio at the University of Alberta for providing the pJW15 (pP*_wec_*::*lux* reporter) plasmid, and Dr. Renato Morona at the University of Adelaide for the ECA antibody. This work was funded by the National Institute of Allergy and Infectious Diseases under Award Numbers R01-AI155915, R01-AI155915-S1, R01-AI192423, and R03-AI177677 (to A.M.M.). The funders played no role in the study design, data collection, analysis, decision to publish, or preparation of the manuscript.

## Notes

### Competing Interest Statement

The authors have declared no competing interest.

### Summary of Updates

Supplementary information is being added to the preprint.

