## Supplemental Material for "Stress-response regulators tune enterobacterial common antigen synthesis to support acid and reactive nitrogen species stress resistance"

#### Supplemental Figures

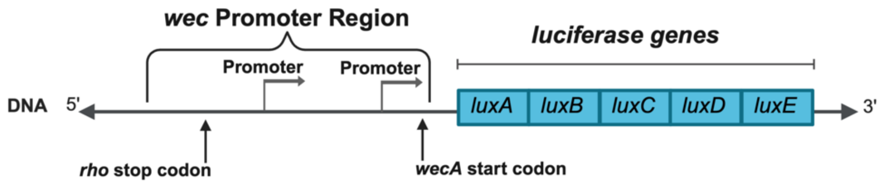


**Fig. S1. Map of transcriptional reporter.** Map depicts a section of the pJW15-Pwec reporter plasmid which includes a 520-base pair *wec* promoter region cloned upstream of the luminescent luciferase genes (blue). The approximate locations of the two predicted *wec* transcriptional start sites are annotated within the promoter region. Created in BioRender. Hill, H. (2026) https://BioRender.com/vupfwm4

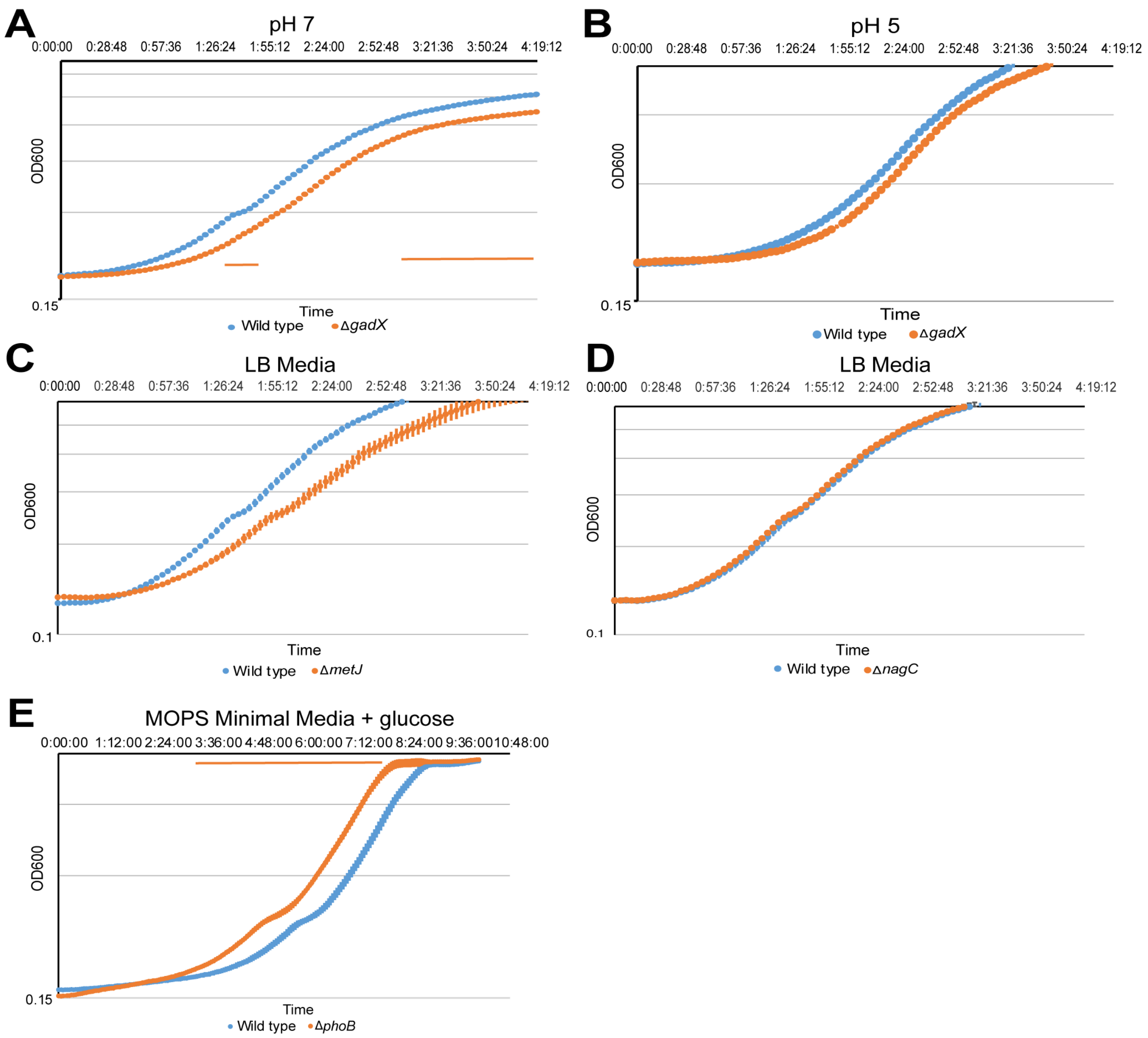


**Fig. S2. Growth rate of mutants is comparable to wild type.** Strains were grown overnight at 37°C in neutral LB media, then back-diluted 1:100 in media of the indicated type and $OD_{600}$ and luminescence were measured. **A.** A Δ*gadX* mutant had a statistically significant decrease in growth in pH 7 media**. B.** A Δ*gadX* mutant grown in pH 5 LB media had no difference in growth compared to the wild type. **C.** A Δ*metJ* mutant grown in LB media had no statistically significant difference in growth vs the wild type. **D.** Growth of a Δ*nagC* mutant in LB media overlaps with the wild type strain **E.** A Δ*phoB* mutant grown in MOPS minimal media with 0.2% glucose had a statistically significant increase in growth compared to the wild type. Bar: $p<0.05$ by Student's T-test.

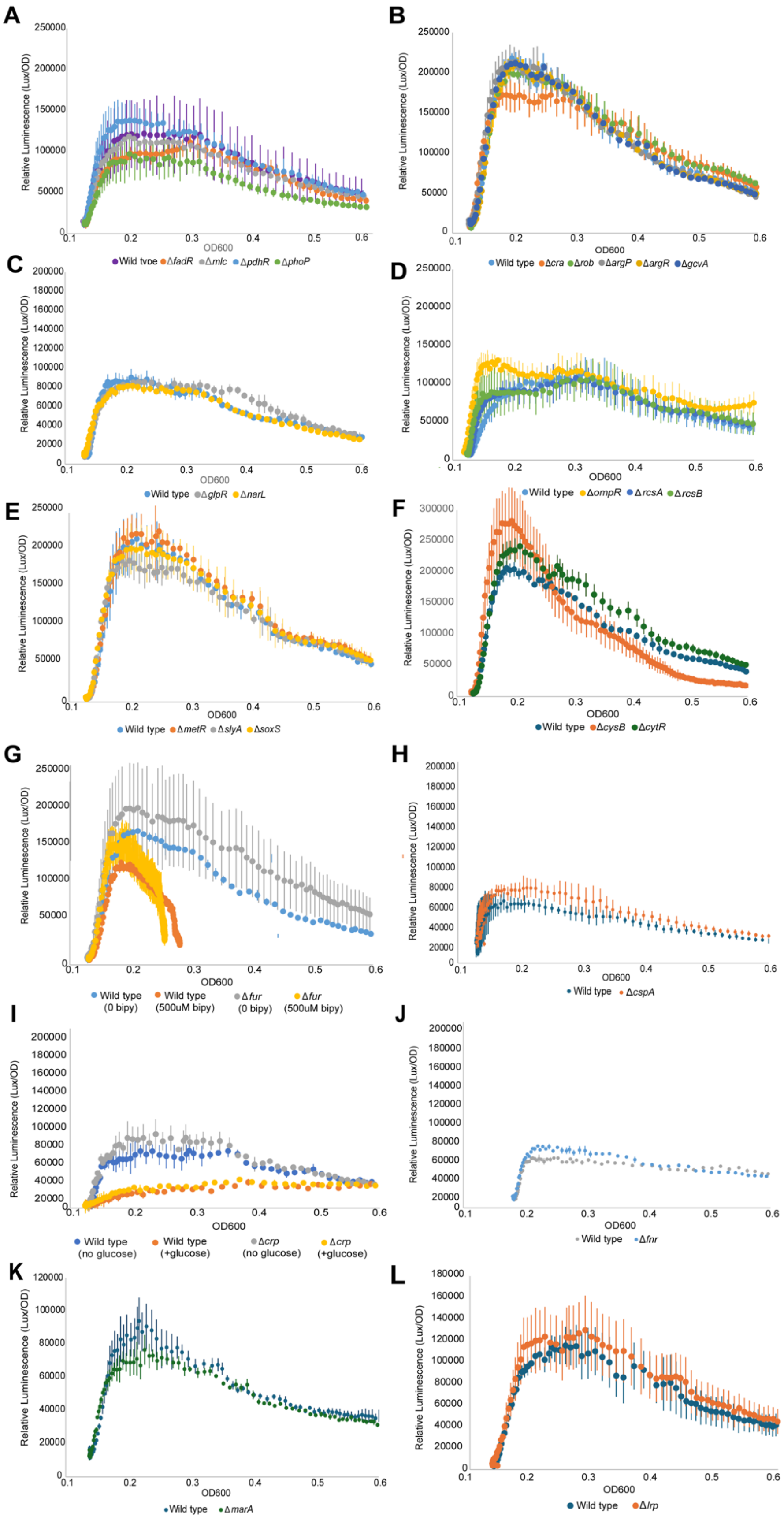
A
Relative Luminescence (Lux/OD)
OD600
Wild type ΔfadR Δmlc ΔpdhR ΔphoP
B
Relative Luminescence (Lux/OD)
OD600
Wild type Δcra Δrob ΔargP ΔargR ΔgcvA
C
Relative Luminescence (Lux/OD)
OD600
Wild type ΔglpR ΔnarL
D
Relative Luminescence (Lux/OD)
OD600
Wild type ΔompR ΔrcsA ΔrcsB
E
Relative Luminescence (Lux/OD)
OD600
Wild type ΔmetR ΔslyA ΔsoxS
F
Relative Luminescence (Lux/OD)
OD600
Wild type ΔcysB ΔcytR
G
Relative Luminescence (Lux/OD)
OD600
Wild type (0 bipy)
Wild type (500uM bipy)
Δfur (0 bipy)
Δfur (500uM bipy)
H
Relative Luminescence (Lux/OD)
OD600
Wild type ΔcspA
I
Relative Luminescence (Lux/OD)
OD600
Wild type (no glucose)
Wild type (+glucose)
Δcrp (no glucose)
Δcrp (+glucose)
J
Relative Luminescence (Lux/OD)
OD600
Wild type Δfnr
K
Relative Luminescence (Lux/OD)
OD600
Wild type ΔmarA
L
Relative Luminescence (Lux/OD)
OD600
Wild type Δlrp

**Fig. S3. Transcriptional reporter assays of potential *wec* operon regulators.** Reporter assays were performed in LB at 37°C, unless otherwise noted. None of the indicated gene deletions caused a significant change in $P_{wec}$ activity from that of wild type. **A.** Deletions of *phoP*, *pdhR*, *mlc*, and *fadR* are shown. **B.** Deletions of *cra*, *rob*, *argP*, *argR*, and *gcvA* are shown. **C.** Deletion of *glpR* and *narL* are shown. **D.** Deletions of *ompR*, *rcsA*, and *rcsB* are shown. **E.** Deletions of *metR*, *slyA*, and *soxS* are shown. **F.** Deletions of *cytR* and *cysB* are shown. **G.** $P_{wec}$ activity in a Δ*fur* strain was tested in LB media and LB with 500 µM 2'2-bipyridil (an iron chelator). **H.** Deletion of *cspA* grown in LB media at 23°C is shown. **I.** $P_{wec}$ activity in a Δ*crp* was tested in LB media and LB with 0.2% glucose. **J.** Deletion of *fnr* is shown. **K.** Deletion of *marA* is shown **L.** Deletion of *lrp* is shown. ArcA was excluded as it is active in anaerobic conditions and the luciferase reporter requires oxygen to function. All data are shown as the mean of three biological replicates ± the SEM. Significance testing was performed by Student's T-test.

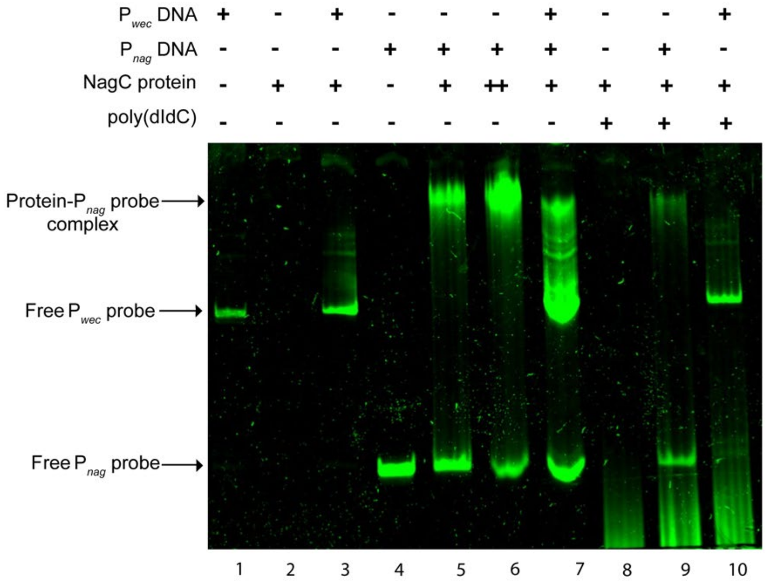


**Fig. S4. NagC does not bind directly the *wec* promoter region**. Experiment performed as shown in Figure 5. Incubation of the $P_{wec}$ DNA probe with NagC did not result in a mobility shift (Lane 3), whereas NagC binding to the positive control $P_{nag}$ DNA resulted in a mobility shift maintained in the presence of competitors (Lanes 5-7, 9). For NagC protein, + represents 200 nM while ++ represents 300 nM. Image is representative of three independent experiments.

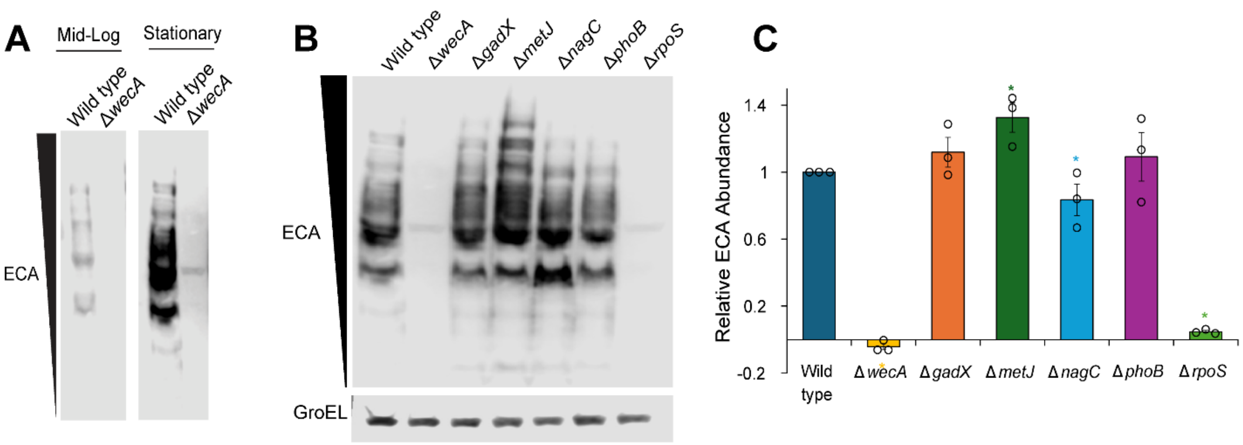


**Fig S5. Transcriptional regulators cause smaller changes in ECA levels in stationary phase. A.** Representative ECA immunoblot showing ECA levels from mid-log (Fig. 4A) and stationary phase (Fig. S4B) samples at the same exposure. Stationary phase samples have higher levels of ECA than sample taken from log phase. **B.** Representative ECA immunoblot conducted with samples taken from overnight cultures. **C.** The identified transcriptional regulators cause smaller changes in ECA levels at this growth stage than during exponential growth. Data are shown as the mean fold ECA signal to wild type of three biological replicates ± the SEM with individual data points. * $p<0.05$ by Wilcoxon test with normal approximation

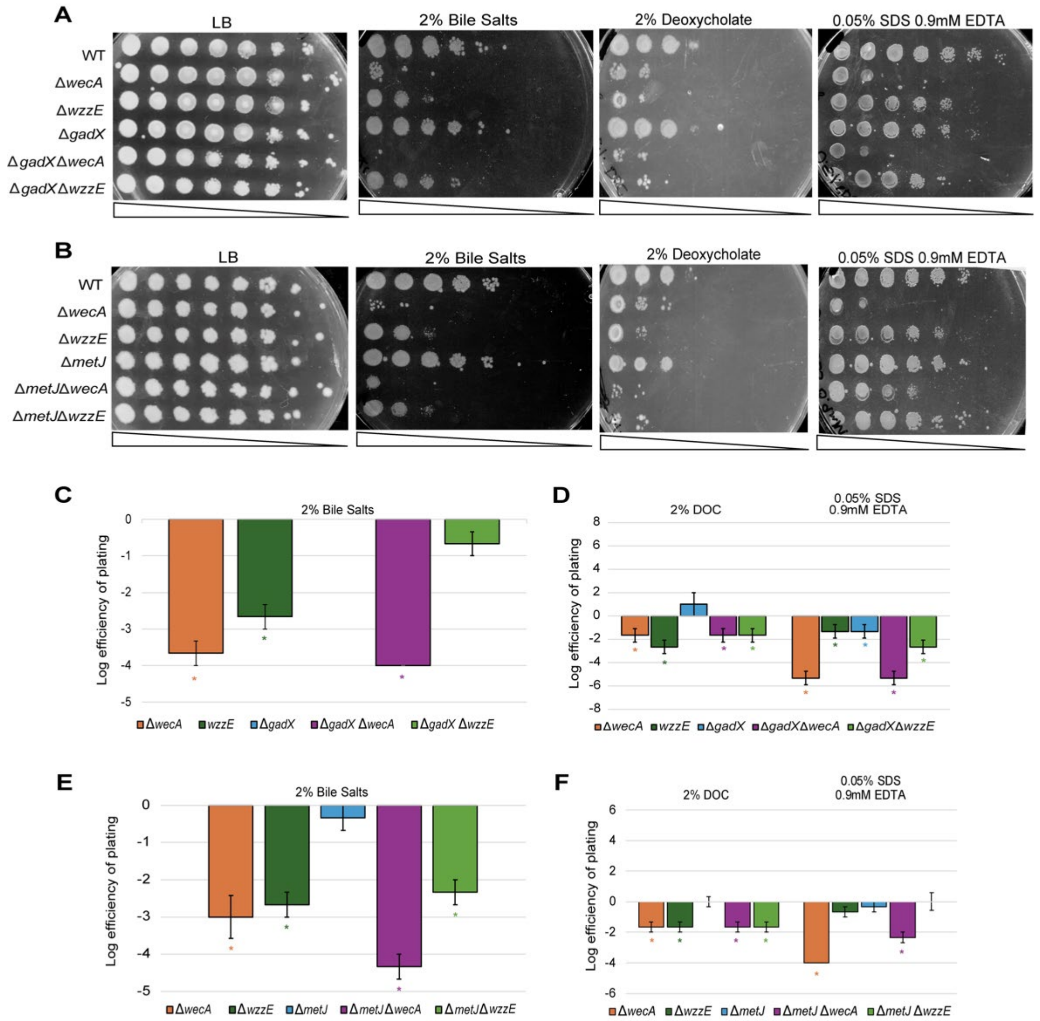


**Fig. S5. Transcriptional regulators and ECA mutants have minimal genetic interactions affecting envelope permeability in stationary phase. A.** Cultures were grown to mid-log phase ($OD_{600}$ ~0.5) then serial diluted into fresh LB, plated on media with 2% bile salts, and grown for 16 hours at 37 °C. Stationary phase cells were plated on either 2% deoxycholate or 0.5% SDS 0.9 mM EDTA. Images are representative of three independent experiments. **B.** Cultures were grown to mid-log phase ($OD_{600}$ ~0.5) then serial diluted into fresh LB, plated on media with 2% bile salts and grown for 16 hours at 37 °C. Stationary phase cells were plated on either 2% deoxycholate or 0.5% SDS 0.9 mM EDTA. **C.** Mid-log cells with Δ*gadX* in conjunction with Δ*wzzE* or Δ*wecA* had no significant change in bile salt sensitivity than their respective parent strains. **D.** EOPs were performed with stationary phase cultures and strains with Δ*gadX* in combination with *wec* mutations did not have a significant change in sensitivity to deoxycholate or SDS-EDTA compared to *wec* mutants do alone. **E.** The Δ*metJ* Δ*wecA* or Δ*metJ* Δ*wzzE* mid-log cells did not have a significant change in sensitivity to bile salts compared to their parent strains. **F.** Stationary phase Δ*metJ* Δ*wzzE* and Δ*metJ* Δ*wecA* cells show no significant change in deoxycholate or SDS-EDTA sensitivity compared to the single deletions when plated from stationary phase . There was no significant difference between the *gadX* or *metJ* double mutants and their parent strain by Mann-Whitney test with Benjamani-Hochberg (FDR) correction. Data are shown as the mean CFU log10 fold to wild type of three biological replicates ± the SEM,* p<0.05 comparison of each strain to wild type by one-way ANOVA with Bonferroni correction.

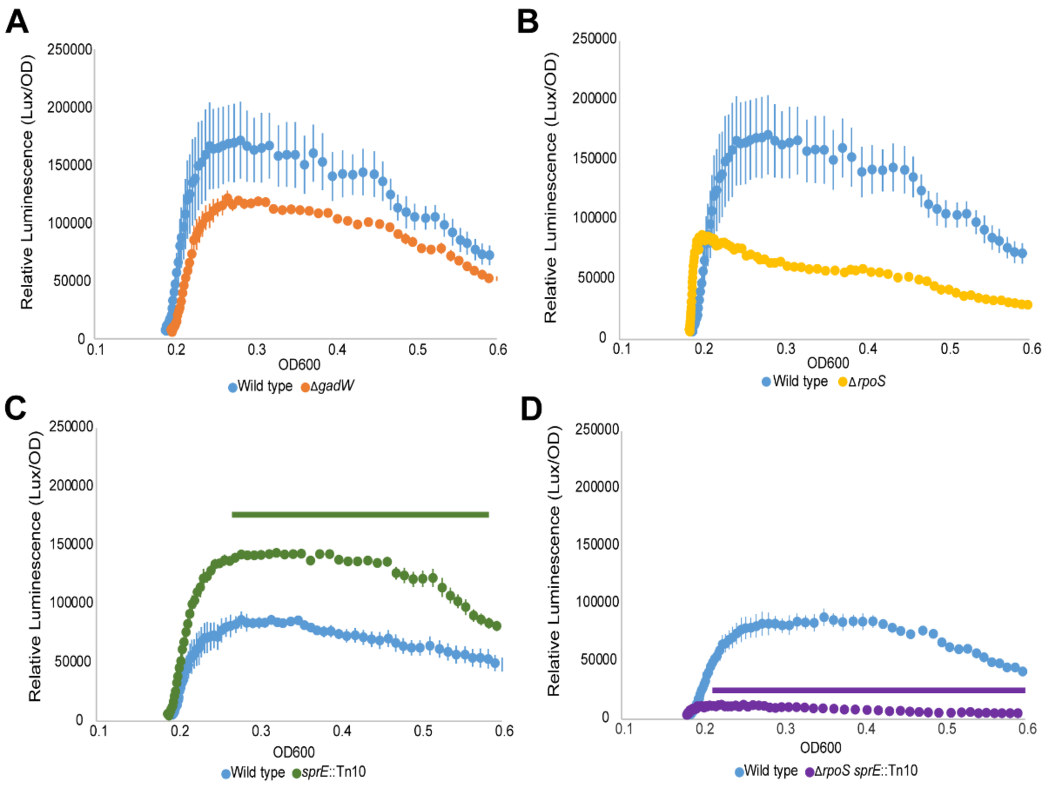


**Fig. S7. RpoS positively regulates P$_{wec}$ activity.** Strains were grown overnight in neutral pH LB media, then back-diluted 1:100 in mildly acidic pH (5.0) LB media for P$_{wec}$ reporter assays. **A.** Δ*gad*W does not change P$_{wec}$ activity. **B.** Δ*rpoS* causes a non-significant decreased P$_{wec}$ activity, suggesting RpoS might positively regulate P$_{wec}$. Wild-type, Δ*gad*W, and Δ*rpoS* were run simultaneously and wild-type data from these experiments are shown in (A) and (B). **C.** In contrast, *sprE*::Tn*10*, which increases RpoS levels, increases P$_{wec}$ activity significantly compared to the wild type. **D.** Δ*rpoS* is epistatic to *sprE*::Tn*10* demonstrating SprE acts on P$_{wec}$ activity through RpoS. Data are shown as the mean of three biological replicates ± the SEM. Significance was determined by Student's T-test.

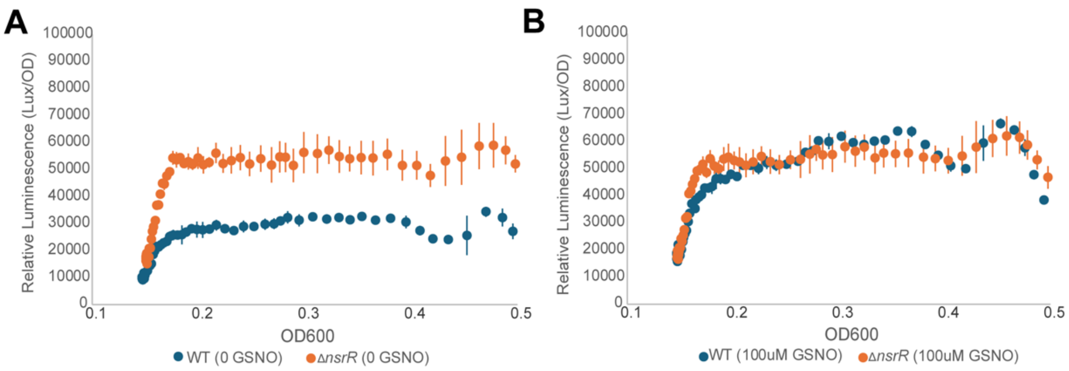


**Fig. S8. Deletion of *nsrR* has a non-significant effect on P*wec* activity that is alleviated by addition of GSNO.** Strains were grown overnight in pH LB media, then back-diluted 1:100 in fresh LB media and grown kinetically at 37°C for $P_{wec}$ reporter assays **A.** Transcriptional reporter assay shows a non-significant increase $P_{wec}$ activity in the vehicle-treated Δ*nsrR* cells compared to vehicle-treated wild type cells**. B.** Wild type $P_{wec}$ activity overlaps with that of the Δ*nsrR* mutant in the presence of 100 μM of GSNO.

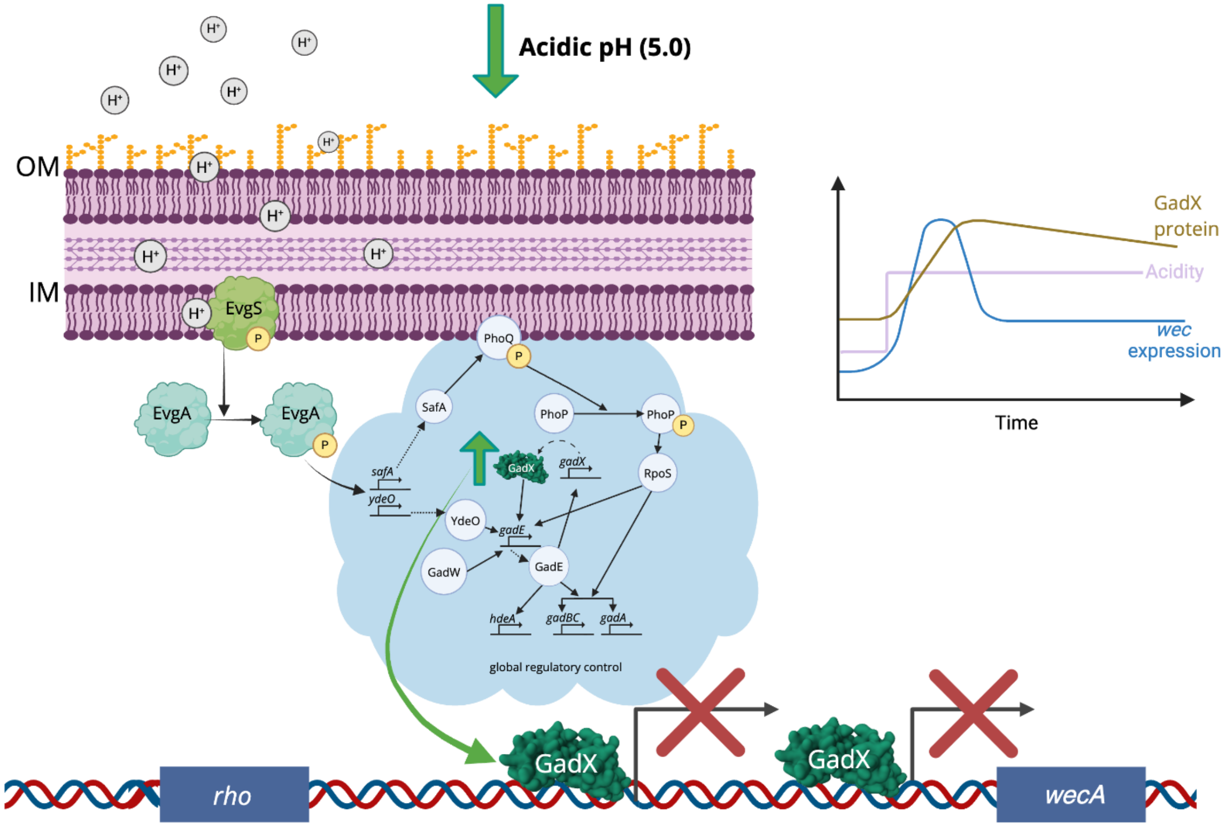


**Fig. S9. Model for *wec* transcriptional regulation by GadX.** In the glutamate-dependent acid stress response system, increased hydrogen ions in the periplasm are sensed by the sensor kinase EvgS, which interacts with and phosphorylates EvgA. Phosphorylated EvgA then activates transcription of acid stress response regulators *safA* and *ydeO*. SafA phosphorylates PhoQ, which then phosphorylates PhoP, which activates RpoS. RpoS activates transcription of *gadE,* the master acid response regulator, *gadBC,* and *gadA.* GadW and YdeO also act directly on *gadE* transcription. GadE activates transcription of *hdeA* and *gadX* and GadX also autoregulates its own expression as well as affecting the transcription of *gadE*. In our model, GadX binds to repress transcription of the *wec* operon at neutral pH; however, we hypothesize that, when the cell experiences acid stress, an acid stress regulator(s), possibly RpoS, increases *wec* operon expression to help the cell maintain efficient growth. GadX levels then increase and more GadX binds to the *wec* promoters to turn down ECA production, leading to moderated *wec* operon expression under sustained acid exposure. Created in BioRender. Hill, H. (2026) https://BioRender.com/tb6ue87

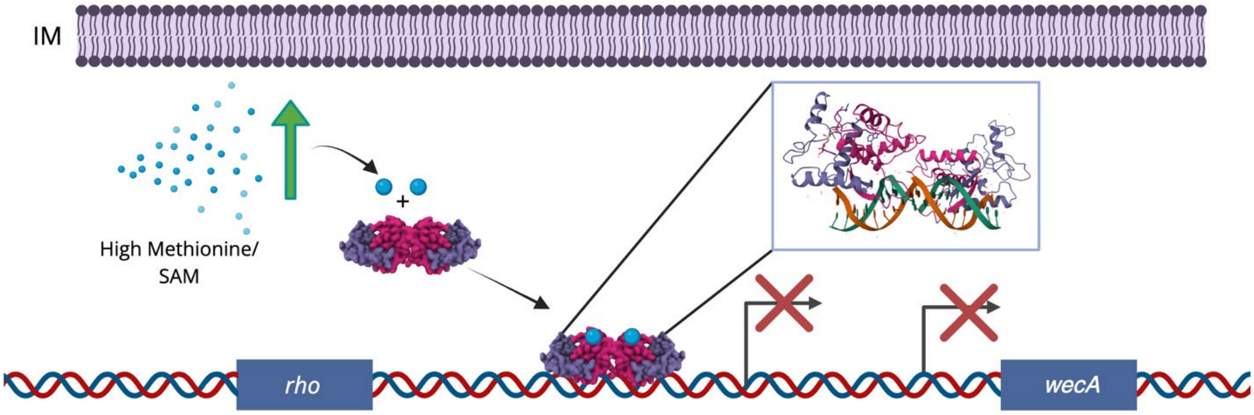


**Fig. S10. Model for *wec* transcriptional regulation by MetJ.** In our model, when methionine and S-adenosylmethionine levels are high, SAM binds the MetJ homodimer which then binds operators upstream of *wecA* to repress *wec* transcription and decrease production of ECA. However, ECA production is derepressed in the presence of GSNO or other reactive nitrogen species, which decrease methionine biosynthesis and induce stress responses. Created in BioRender. Hill, H. (2026) https://BioRender.com/6sw9cgk

### Supplemental Tables

#### Table S1. Potential *wec* operon regulators[1].

| PWM (Species)[2] | Start Position[3] | End Position[3] | Strand | Score[4] | Max Score[4] | Difference[4] | Sequence[5] |
|---|---|---|---|---|---|---|---|
| OxyR (SELEX) \| Escherichia coli (strain K12) | 394 | 439 | - | 13.54 | 22.11 | 8.57 | GGGAAAGTCCGTTCCAAACCCACCAGTCCTGTGCAAAAATTAACCA |
| | 230 | 275 | + | 13.27 | 22.11 | 8.84 | ACGATTTCTTCGAAATGATGAAACGCTCATAAATTTGTCTTATGCC |
| | 429 | 474 | - | 12.87 | 22.11 | 9.24 | AGCAGAAGTATAACCACGAAGACCTTTATTCAGAAGGGAAAGTCCG |
| CysB \| Escherichia coli (strain K12) | 365 | 395 | - | 9.36 | 11.43 | 2.07 | CATTTGATATGCAAATTGCATAACATGCCAG |
| ArgR \| Escherichia coli (strain K12) | 151 | 164 | - | 8.84 | 10.35 | 1.51 | TGAATGATTTTGCG |
| | 366 | 379 | - | 8.25 | 10.35 | 2.1 | TGCATAACATGCCA |
| | 443 | 456 | + | 8.11 | 10.35 | 2.24 | TGAATAAAGGTCTT |
| CspA \| Escherichia coli (strain K12) | 224 | 228 | + | 10 | 10 | 0 | CCAAT |
| GcvA \| Escherichia coli (strain K12) | 473 | 477 | + | 10 | 10 | 0 | CTAAT |
| MetR \| Escherichia coli (strain K12) | 51 | 57 | + | 9.03 | 9.5 | 0.47 | TGAAAAA |
| | 443 | 449 | + | 8.74 | 9.5 | 0.76 | TGAATAA |
| Fnr \| Escherichia coli (strain K12) | 214 | 227 | - | 6.74 | 9.01 | 2.27 | TTGGTCTTGGTCAT |
| | 142 | 155 | - | 6.35 | 9.01 | 2.66 | TTGCGCAGGATCCA |
| | 264 | 277 | - | 6.32 | 9.01 | 2.69 | TTGGCATAAGACAA |
| Crp \| Escherichia coli (strain K12) | 392 | 413 | - | 6.25 | 8.83 | 2.58 | TCCTGTGCAAAAATTAACCATT |
| | 148 | 169 | + | 5.92 | 8.83 | 2.91 | CTGCGCAAAATCATTCACCCGA |
| | 482 | 503 | - | 5.87 | 8.83 | 2.96 | CACAATGCATGCTCTCAGAGAA |
| OmpR \| Escherichia coli (strain K12) | 52 | 58 | + | 8.25 | 8.65 | 0.4 | GAAAAAC |
| | 478 | 484 | - | 8.13 | 8.65 | 0.52 | GAAAATT |
| | 347 | 353 | + | 8.13 | 8.65 | 0.52 | GAAAATT |
| CytR \| Escherichia coli (strain K12) | 375 | 386 | - | 7.78 | 8.08 | 0.3 | TGCAAATTGCAT |
| | 474 | 485 | - | 7.65 | 8.08 | 0.43 | AGAAAATTATTA |
| | 134 | 145 | + | 7.65 | 8.08 | 0.43 | AGAAAATGTGGA |
| GlpR \| Escherichia coli (strain K12) | 147 | 166 | + | 6.29 | 7.8 | 1.51 | CCTGCGCAAAATCATTCACC |
| | 413 | 432 | - | 6.2 | 7.8 | 1.6 | TCCGTTCCAAACCCACCAGT |
| | 168 | 187 | + | 6.16 | 7.8 | 1.64 | GATGGGCGAAATCGATGCAA |
| IHF \| Escherichia coli (strain K12) | 387 | 402 | + | 6.41 | 7.76 | 1.35 | TATCAAATGGTTAATT |
| | 198 | 213 | - | 5.72 | 7.76 | 2.04 | TGCCAGTTTATTAATG |
| | 355 | 370 | - | 5.57 | 7.76 | 2.19 | TGCCAGCAAATTCCAG |
| | 237 | 245 | + | 6.82 | 7.58 | 0.76 | CTTCGAAAT |
| | 421 | 429 | - | 6.59 | 7.58 | 0.99 | GTTCCAAAC |

| PWM (Species)[2] | Start Position[3] | End Position[3] | Strand | Score[4] | Max Score[4] | Difference[4] | Sequence[5] |
|---|---|---|---|---|---|---|---|
| MetJ \| Escherichia coli (strain K12) | 3 | 11 | - | 5.9 | 7.58 | 1.68 | CCTTTAAAC |
| GadX \| Escherichia coli (strain K12) | 295 | 312 | + | 5.81 | 7.55 | 1.74 | GTGGCGTTTTGCTTTTAT |
| | 448 | 465 | + | 5.53 | 7.55 | 2.02 | AAAGGTCTTCGTGGTTAT |
| | 182 | 199 | - | 5.47 | 7.55 | 2.08 | TGAGGAATTCCATTGCAT |
| ArcA \| Escherichia coli (strain K12) | 341 | 350 | + | 6.11 | 7.5 | 1.39 | TGTTAGGAAA |
| | 395 | 404 | + | 6.1 | 7.5 | 1.4 | GGTTAATTTT |
| | 371 | 380 | + | 6.06 | 7.5 | 1.44 | TGTTATGCAA |
| PdhR \| Escherichia coli (strain K12) | 389 | 394 | + | 6.31 | 6.93 | 0.62 | TCAAAT |
| | 224 | 229 | + | 6.27 | 6.93 | 0.66 | CCAATG |
| H-NS \| Escherichia coli (strain K12) | 179 | 188 | + | 5.4 | 6.75 | 1.35 | TCGATGCAAT |
| | 246 | 255 | - | 5.26 | 6.75 | 1.49 | GCGTTTCATC |
| | 255 | 264 | + | 5.17 | 6.75 | 1.58 | CTCATAAATT |
| Mlc \| Escherichia coli (strain K12) | 240 | 245 | + | 6.15 | 6.44 | 0.29 | CGAAAT |
| | 174 | 179 | + | 6.15 | 6.44 | 0.29 | CGAAAT |
| | 151 | 156 | + | 5.7 | 6.44 | 0.74 | CGCAAA |
| FadR \| Escherichia coli (strain K12) | 412 | 418 | + | 5.8 | 6.44 | 0.64 | GACTGGT |
| | 87 | 93 | + | 5.54 | 6.44 | 0.9 | TTCTGGT |
| | 206 | 212 | + | 5.51 | 6.44 | 0.93 | AACTGGC |
| RcsAB \| Escherichia coli (strain K12) | 344 | 351 | + | 5.65 | 5.93 | 0.28 | TAGGAAAA |
| Lrp \| Escherichia coli (strain K12) | 442 | 456 | - | 5.02 | 5.86 | 0.84 | AAGACCTTTATTCAG |
| | 366 | 380 | + | 4.64 | 5.86 | 1.22 | TGGCATGTTATGCAA |
| | 199 | 213 | - | 4.56 | 5.86 | 1.3 | TGCCAGTTTATTAAT |
| ArgP \| Escherichia coli (strain K12) | 231 | 242 | + | 4.99 | 5.77 | 0.78 | CGATTTCTTCGA |
| | 380 | 391 | - | 4.91 | 5.77 | 0.86 | TGATATGCAAAT |
| | 312 | 323 | - | 4.86 | 5.77 | 0.91 | AGATTACAGATA |
| GlnG \| Escherichia coli (strain K12) | 327 | 332 | + | 5.46 | 5.23 | -0.23 | TGCCGC |
| | 90 | 95 | + | 5.14 | 5.23 | 0.09 | TGGTAC |
| | 296 | 301 | + | 5.08 | 5.23 | 0.15 | TGGCGT |
| NarL \| Escherichia coli (strain K12) | 465 | 471 | + | 4.06 | 4.3 | 0.24 | TACTTCT |
| | 7 | 13 | - | 4.01 | 4.3 | 0.29 | TACCTTT |
| | 93 | 99 | + | 3.99 | 4.3 | 0.31 | TACCCGT |
| OxyR \| Escherichia coli (strain K12) | 270 | 280 | + | 3.35 | 4.07 | 0.72 | TATGCCAAAAA |
| | 271 | 281 | + | 3.18 | 4.07 | 0.89 | ATGCCAAAAAC |
| | 300 | 310 | - | 3.11 | 4.07 | 0.96 | AAAAGCAAAAC |
| Fis \| Escherichia coli (strain K12) | 395 | 409 | - | 3.35 | 3.6 | 0.25 | GTGCAAAAATTAACC |
| | 472 | 486 | - | 3.2 | 3.6 | 0.4 | GAGAAAATTATTAGC |
| | 364 | 378 | - | 3.19 | 3.6 | 0.41 | GCATAACATGCCAGC |
| Fur (8mer)\| Escherichia coli (strain K12) | 51 | 58 | + | 2.2 | 2.47 | 0.27 | TGAAAAAC |
| | 459 | 466 | + | 2.15 | 2.47 | 0.32 | TGGTTATA |
| | 443 | 450 | + | 2.15 | 2.47 | 0.32 | TGAATAAA |

[1] Analysis conducted with the 2018 version of PRODORIC2 Virtual Footprint tool (68)

[2] Position weight matrix (PWM) for indicated regulator in the indicated species

[3] Positions are given from the start of a 500 bp region upstream of the *wecA* start codon.

[4] The score of the predicted binding site, the maximum possible score for the PWM, and the differences between the scores are indicated.

[5] Sequence of the predicted binding site

**Table S2. Updated list of potential *wec* operon regulators[1]**

| PRODORIC Matrix[2] | Core Score[3] | Rel Score[3] | Binding Site Sequence | Function |
|---|---|---|---|---|
| ArgP | 5.50 | 0.99 | AATTATTA | Arginine metabolism |
| | 5.42 | 0.97 | GTTTATTA | |
| ArgR | 7.82 | 0.79 | CGCAAAATCATTCA | Arginine metabolism |
| | 7.72 | 0.78 | TGAATGATTTTGCG | |
| | 7.68 | 0.77 | GGAATTCCATTGCA | |
| | 7.60 | 0.77 | TGTCAGTAAATTCA | |
| Crp | 5.39 | 0.89 | TGTCAGTAAATTCACA | Central metabolism |
| Cra | 11.50 | 0.91 | TGAAACGCTCAT | Central metabolism |
| CysB | 6.34 | 0.95 | TCATTGCCAGTTTAT | Cysteine metabolism |
| CytR | 4.07 | 1.0 | TGCAA | Nucleoside metabolism |
| Fur | 7.42 | 1.0 | TGCAAT | Iron homeostasis |
| GadX | 7.55 | 0.81 | GTGGCGTTTTGCTTTTAT | Acid stress response |
| | 7.55 | 0.78 | AAAGGTCTTCGTGGTTAT | |
| | 7.55 | 0.74 | TGAGGAATTCCATTGCAT | |
| Lrp | 2.14 | 0.93 | GAGAAAATTATT | Global regulator, entry to stationary phase |
| | 2.03 | 0.88 | CATAAATTTGTC | |
| | 1.99 | 0.87 | AAGACCTTTATT | |
| MarA | 6.31 | 0.87 | AGAGCATGCATTGTGAATTT | Multidrug resistance, redox and envelope stress response, and biofilm formation |
| | 6.19 | 0.85 | ATTGCATAACATGCCAGCAA | |
| MetJ | 5.35 | 0.87 | TAACATCG | Methionine metabolism |
| | 5.18 | 0.84 | TGACGATT | |
| | 4.94 | 0.80 | AGACAAAT | |
| NarL | 4.10 | 0.97 | TACTTCT | Anaerobic growth, nitrogen metabolism |
| | 4.02 | 0.95 | TACAGAT | |
| | 4.02 | 0.95 | TACCTTT | |
| OmpR | 6.07 | 0.82 | TAAGATTACAGATATAAA | Osmolarity and acid stress |
| | 5.55 | 0.75 | TTAGCAGAAGTATAACCA | |
| | 5.47 | 0.74 | TGGAATTCCTCATTAATA | |
| OxyR | 5.70 | 0.81 | TAAGATTACAGATAT | Oxidative stress regulator |
| | 5.33 | 0.76 | TTTGCTTTTATATCT | |
| | | 4.07 | AAAAGCAAAAC | |
| PhoB | 5.11 | 0.82 | ATGTTATGCAATTTGCATATCA | Phosphate uptake regulator |
| | 4.93 | 0.79 | TTGATATGCAAATTGCATAACA | |
| | 4.92 | 0.79 | ATATCTGTAATCTTAATGCCGC | |
| | 4.90 | 0.78 | GTTTCATCATTTCGAAGAAATC | |

| PRODORIC Matrix[2] | Core Score[3] | Rel Score[3] | Binding Site Sequence | Function |
|---|---|---|---|---|
| | 4.70 | 0.75 | CTCTCGTAAGATCGCTGAAAAA | |
| | 4.66 | 0.75 | ATTTCGCCCATCGGGTGAATGA | |
| | 4.65 | 0.74 | TTTGCTGGCATGTTATGCAATT | |
| | 4.63 | 0.74 | ATCGCTGAAAAACGCGTCTTCC | |
| | 4.52 | 0.72 | GTTTTTGGCATAAGACAAATTT | |
| | 4.47 | 0.72 | TTTAAAGGTACAGGCAACATGG | |
| | 4.44 | 0.71 | CTTACGAGAGAGGTGCAGTTCC | |
| Rob | 9.06 | 0.83 | ATTGCATAACATGCCAGCAA | Antibiotic resistance, stress responses, metabolism, and cell envelope functions |
| RcsAB | 9.60 | 0.95 | TAGGAAAATTCCTG | Envelope stress response |
| SlyA | 7.01 | 0.87 | TTATATCTGTAA | Fimbriae and capsule/hemolysin; Virulence genes |
| SoxS | 5.44 | 0.91 | ATTGCATAACATGCCAGCAA | Oxidative stress response |
| | 5.42 | 0.90 | AGAGCATGCATTGTGAATTT | |
| | 5.17 | 0.86 | CTGGCATGTTATGCAATTTG | |
| | 4.97 | 0.83 | AATGCATGCTCTCAGAGAAA | |
| | 4.96 | 0.83 | TATGCAAATTGCATAACATG | |
| | 4.91 | 0.82 | CCAGCAAATTCCAGGAATTT | |
| | 4.90 | 0.82 | CTGGCAATGACCAAGACCAA | |
| | 4.85 | 0.81 | TGCGCAAAATCATTCACCCG | |
| | 4.80 | 0.80 | ACTGCAGAAAATGTGGATCC | |
| | 4.75 | 0.79 | ATTGCATCGATTTCGCCCAT | |
| | 4.72 | 0.79 | ACTGCACCTCTCTCGTAAGA | |
| | 4.61 | 0.77 | TTTGCACAGGACTGGTGGGT | |
| | 4.59 | 0.76 | TGTGCAAAAATTAACCATTT | |

[1] Analysis conducted with version 1.4 of PRODORIC's Virtual Footprint Promoter Analysis tool (2023) (68)

[2] PWM for indicated regulator in *E. coli* K-12

[3] The score of the predicted binding site and the score relative to the maximum possible score for the PWM are indicated.

**Table S3. Strains and Plasmids used in this study**

| Strain | Genotype | Reference |
|---|---|---|
| MG1655 | K-12 F– λ– *rph-1* | (69) |
| **HB316** | W3110 pCA24N-*gadX* | (67) |
| **HB368** | MG1655 Δ*metJ* pCA24N-*metJ* | This study |
| **HB360** | MG1655 pCA24N-*nagC* | This study |
| **HB317** | W3110 pCA24N-*phoB* | (67) |
| **HB276** | MG1655 Δ*wecA* | This study |
| **HB516** | MG1655 Δ*waaL* | This study |
| **HB293** | MG1655 Δ*wzzE* | This study |
| **HB562** | MG1655 Δ*wzzE* Δ*waaL* | This study |
| **HB194** | MG1655 pJW15-$P_{wec}$ | (67) |
| **HB195** | MG1655 pJW15 no promoter | This study |
| **HB024** | MG1655 Δ*cspA* pJW15-$P_{wec}$ | This study |
| **HB025** | MG1655 Δ*pdeL* pJW15-$P_{wec}$ | This study |
| **HB026** | MG1655 Δ*glnG* pJW15-$P_{wec}$ | This study |
| **HB029** | MG1655 Δ*lrp* pJW15-$P_{wec}$ | This study |
| **HB030** | MG1655 Δ*fadR* pJW15-$P_{wec}$ | This study |
| **HB037** | MG1655 Δ*glpR* pJW15-$P_{wec}$ | This study |
| **HB042** | MG1655 Δ*arcA* pJW15-$P_{wec}$ | This study |
| **HB052** | MG1655 Δ*crp* pJW15-$P_{wec}$ | This study |
| **HB053** | MG1655 Δ*mlc* pJW15-$P_{wec}$ | This study |
| **HB061** | MG1655 Δ*fnr* pJW15-$P_{wec}$ | This study |
| **HB062** | MG1655 Δ*fur* pJW15-$P_{wec}$ | This study |
| **HB068** | MG1655 Δ*pdhR* pJW15-$P_{wec}$ | This study |
| **HB069** | MG1655 Δo*xyR* pJW15-$P_{wec}$ | This study |
| **HB074** | MG1655 Δ*cysB* pJW15-$P_{wec}$ | This study |
| **HB090** | MG1655 Δ*argR*- pJW15-$P_{wec}$ | This study |
| **HB096** | MG1655 Δ*ihfA* pJW15-$P_{wec}$ | This study |
| **HB097** | MG1655 Δ*argP* pJW15-$P_{wec}$ | This study |
| **HB098** | MG1655 Δ*gcvA* pJW15-$P_{wec}$ | This study |
| **HB112** | MG1655 Δ*metR* pJW15-$P_{wec}$ | This study |
| **HB116** | MG1655 Δ*soxS* pJW15-$P_{wec}$ | This study |
| **HB117** | MG1655 Δ*slyA* pJW15-$P_{wec}$ | This study |
| **HB123** | MG1655 Δ*ihfB* pJW15-$P_{wec}$ | This study |
| **HB129** | MG1655 Δ*cra* pJW15-$P_{wec}$ | This study |
| **HB130** | MG1655 Δ*rob* pJW15-$P_{wec}$ | This study |
| **HB132** | MG1655 Δ*nagC* pJW15-$P_{wec}$ | This study |
| **HB134** | MG1655 Δ*metJ* pJW15-$P_{wec}$ | This study |

| | | |
|---|---|---|
| **HB135** | MG1655 Δ*marA* pJW15-$P_{wec}$ | This study |
| **HB158** | MG1655 Δ*gadW* pJW15-$P_{wec}$ | This study |
| **HB167** | MG1655 Δ*rssB* pJW15-$P_{wec}$ | This study |
| **HB170** | MG1655 Δ*phoP* ΔgadX pJW15-$P_{wec}$ | This study |
| **HB256** | MG1655 Δ*phoB* pJW15-$P_{wec}$ | This study |
| **HB210** | MG1655 Δ*metJ* pJW15-$P_{wec}$ | This study |
| **HB211** | MG1655 Δ*nagC* pJW15-$P_{wec}$ | This study |
| **HB260** | MG1655 Δ*gadX* pJW15-$P_{wec}$ | This study |
| **HB240** | MG1655 Δ*narL* pJW15-$P_{wec}$ | This study |
| **HB247** | MG1655 Δ*gadX* Δ*wzzE* | This study |
| **HB250** | MG1655 Δ*metJ* Δ*wzzE* | This study |
| **HB261** | MG1655 Δ*metJ* Δ*wecA* | This study |
| **HB263** | MG1655 Δ*gadX* Δ*wecA* | This study |
| **HB319** | MG1655 Δ*fnr* pJW15-$P_{wec}$ | This study |
| **HB320** | MG1655 Δ*fadR* pJW15-$P_{wec}$ | This study |
| **HB330** | MG1655 Δ*phoP* pJW15-$P_{wec}$ | This study |
| **HB371** | MG1655 Δ*nsrR* pJW15-$P_{wec}$ | This study |
| **HB372** | MG1655 Δ*abgR* pJW15-$P_{wec}$ | This study |
| **HB547** | MG1655 Δ*rpoS* pJW15-$P_{wec}$ | This study |
